# Bayesian estimation of gene constraint from an evolutionary model with gene features

**DOI:** 10.1101/2023.05.19.541520

**Authors:** Tony Zeng, Jeffrey P. Spence, Hakhamanesh Mostafavi, Jonathan K. Pritchard

## Abstract

Measures of selective constraint on genes have been used for many applications including clinical interpretation of rare coding variants, disease gene discovery, and studies of genome evolution. However, widely-used metrics are severely underpowered at detecting constraint for the shortest ~25% of genes, potentially causing important pathogenic mutations to be overlooked. We developed a framework combining a population genetics model with machine learning on gene features to enable accurate inference of an interpretable constraint metric, *s*_het_. Our estimates outperform existing metrics for prioritizing genes important for cell essentiality, human disease, and other phenotypes, especially for short genes. Our new estimates of selective constraint should have wide utility for characterizing genes relevant to human disease. Finally, our inference framework, GeneBayes, provides a flexible platform that can improve estimation of many gene-level properties, such as rare variant burden or gene expression differences.

## 1 Introduction

Identifying the genes important for disease and fitness is a central goal in human genetics. One particularly useful measure of importance is how much natural selection constrains a gene [1–4]. Constraint has been used to prioritize *de novo* and rare variants for clinical followup [5, 6], predict the toxicity of drugs [7], link GWAS hits to genes [8], and characterize transcriptional regulation [9, 10], among many other applications.

To estimate the amount of constraint on a gene, several metrics have been developed using loss-of-function variants (LOFs), such as protein truncating or splice disrupting variants. If a gene is important, then natural selection will act to remove LOFs from the population. Several metrics of gene importance have been developed based on this intuition to take advantage of large exome sequencing studies.

In one line of research, the number of observed unique LOFs is compared to the expected number under a model of no selective constraint. This approach has led to the widely-used metrics pLI [11] and LOEUF [12].

While pLI and LOEUF have proved useful for identifying genes intolerant to LOF mutations, they have important limitations [3]. First, they are uninterpretable in that they are only loosely related to the fitness consequences of LOFs. Their relationship with natural selection depends on the study’s sample size and other technical factors [3]. Second, they are not based on an explicit population genetics model so it is impossible to compare a given value of pLI or LOEUF to the strength of selection estimated for variants other than LOFs [3, 4].

Another line of research has solved these issues of interpretability by estimating the fitness reduction for heterozygous carriers of a LOF in any given gene [1,2,4]. Throughout, we will adopt the notation of Cassa and colleagues and refer to this reduction in fitness as *s*_het_ [1, 2], although the same population genetic quantity has been referred to as *hs* [4, 13]. In [1], a deterministic approximation was used to estimate *s*_het_, which was relaxed to incorporate the effects of genetic drift in [2]. This model was subsequently extended by Agarwal and colleagues to include the X chromosome and applied to a larger dataset, with a focus on the interpretability of *s*_het_ [4].

A major issue for most previous methods is that thousands of genes have few expected unique LOFs under neutrality, as they have short protein-coding sequences. For example, when LOEUF was introduced [12], it was stated that the method is underpowered for genes with fewer than 10 expected unique LOFs, corresponding to ~25% of genes. This problem is not limited to LOEUF, however, and all of these methods are severely underpowered to detect selection for this ~25% of genes. Throughout, we will say that genes have “few expected LOFs” if they fall in this bottom quartile of genes.

Here, we present an approach that can accurately estimate *s*_het_ even for genes with few expected LOFs, while maintaining the interpretability of previous population-genetics based estimates [1, 2, 4].

Our approach has two main technical innovations. First, we use a novel population genetics model of LOF allele frequencies. Previous methods have either only modeled the number of unique LOFs, throwing away frequency information [11, 12, 14], or considered the sum of LOF frequencies across the gene [1, 2, 4], an approach that is not robust to what we will refer to as misannotated LOFs. In particular, some variants that have been annotated as LOFs do not actually affect the function of a gene product. For example, a splice-disrupting variant may be rescued by a nearby cryptic splice site, or an early stop codon may be in an exon that is absent in physiologically relevant isoforms. In contrast to previous approaches, we model the frequencies of individual LOF variants, allowing us to not only use the information in such frequencies but also to model the possibility that a LOF has been misannotated and hence is expected to evolve neutrally. Our approach uses new computational machinery, described in a companion paper [15], to accurately obtain the likelihood of observing a LOF at a given frequency without resorting to simulation [2, 4] or deterministic approximations [1].

Second, our approach uses thousands of gene features, including gene expression patterns, protein structure information, and evolutionary constraint, to improve estimates for genes with few expected LOFs. By using these features, we can share information across similar genes. Intuitively, this allows us to improve estimates for genes with few expected LOFs by leveraging information from genes with similar features that do have sufficient LOF data.

Adopting a similar approach, a recent paper [14] used gene features in a deep learning model to improve estimation of constraint for genes with few expected LOFs, but did not use an explicit population genetics model, resulting in the same issues with interpretability faced by pLI and LOEUF.

We applied our method to a large exome sequencing cohort [12]. Our estimates of *s*_het_ are substantially more predictive than previous metrics at prioritizing essential and disease-associated genes. We also interrogated the relationship between gene features and natural selection, finding that evolutionary conservation, protein structure, and expression patterns are more predictive of *s*_het_ than co-expression and protein-protein interaction networks. Expression patterns in the brain and expression patterns during development are particularly predictive of *s*_het_. Finally, we use *s*_het_ to highlight differences in selection on different categories of genes and consider *s*_het_ in the context of selection on variants beyond LOFs.

Our approach, GeneBayes, is extremely flexible and can be applied to improve estimation of numerous gene properties beyond *s*_het_. Our implementation is available at https://github.com/tkzeng/GeneBayes.

## 2 Results

### 2.1 Model Overview

Using LOF data to infer gene constraint is challenging for genes with few expected LOFs, with metrics like LOEUF considering almost all such genes to be unconstrained (Figures 1**A,B**). We hypothesized that it would be possible to improve estimation using auxiliary information that may be predictive of LOF constraint, including gene expression patterns across tissues, protein structure, and evolutionary conservation. Intuitively, genes with similar features should have similar levels of constraint. By pooling information across groups of similar genes, constraint estimated for genes with sufficient LOF data may help improve estimation for underpowered genes.

**Figure 1:**
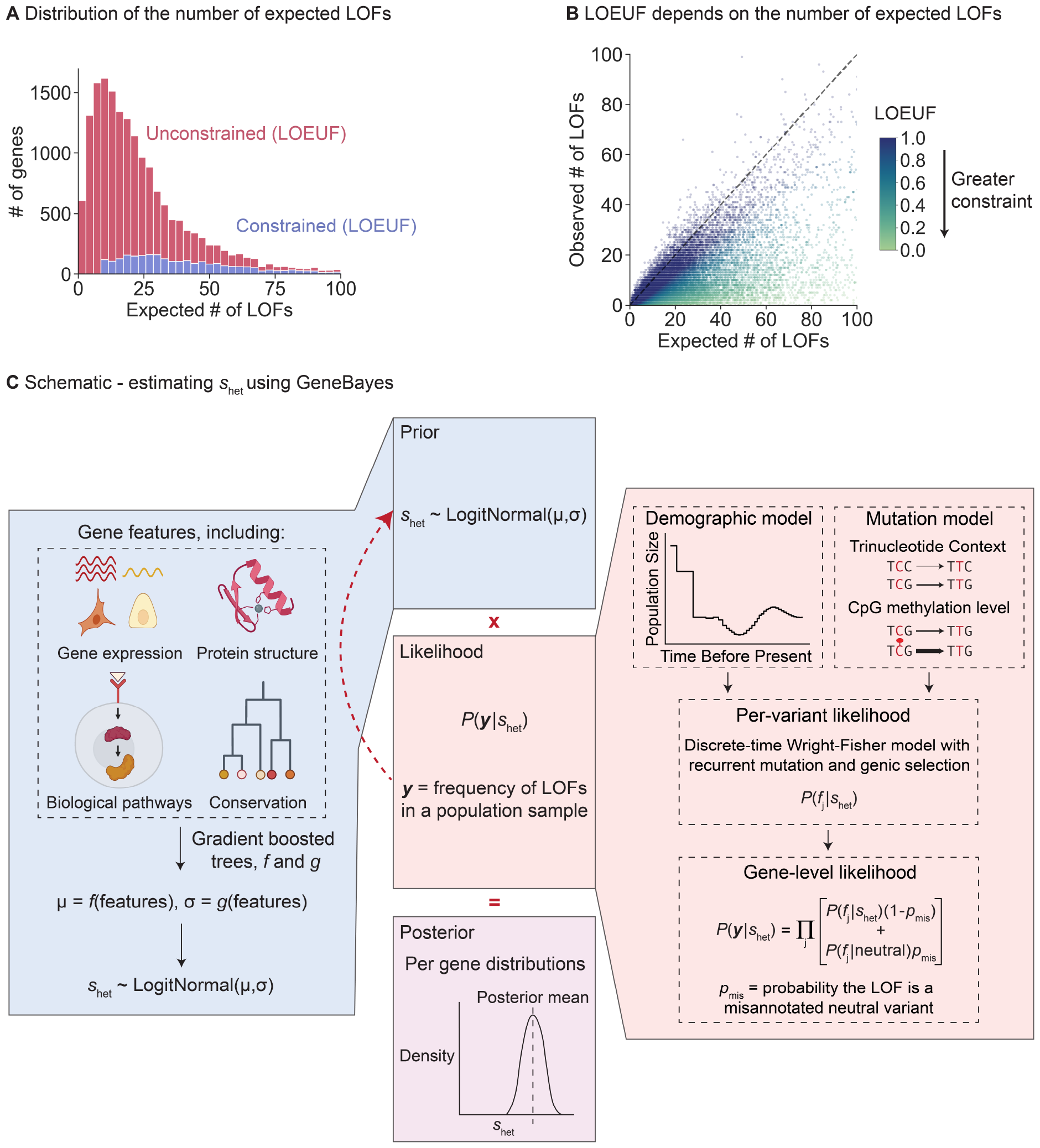
Limitations of LOEUF and schematic for inferring s_het_ using GeneBayes. **A)** Stacked histogram of the expected number of unique LOFs per gene, where the distribution for genes considered unconstrained (respectively constrained) by LOEUF are colored in red (respectively blue). Genes with LOEUF < 0.35 are considered constrained, while all other genes are unconstrained (Methods). The plot is truncated on the x-axis at 100 expected LOFs. **B)** Scatterplot of the observed against the expected number of unique LOFs per gene. The dashed line denotes observed = expected. Each point is a gene, colored by its LOEUF score; genes with LOEUF > 1 are colored as LOEUF = 1. **C)** Schematic for estimating s_het_ using GeneBayes, highlighting the major components of the model: prior (blue boxes) and likelihood (red boxes). Parameters of the prior are learned by maximizing the likelihood (red arrow). Combining the prior and likelihood produces posteriors over s_het_ (purple box). See Methods for details.

However, while the frequencies of LOFs can be related to *s*_het_ through models from population genetics [1, 2, 4], we lack an understanding of how other gene features relate to constraint *a priori*.

To address this problem, we developed a flexible empirical Bayes framework, GeneBayes, that learns the relationship between gene features and *s*_het_ (Figure 1**C**, Methods and Supplementary Note A). Our model consists of two main components. First, we model the prior on *s*_het_ for each gene as a function of its gene features (Figure 1**C**, left). Specifically, we train gradient-boosted trees using a modified version of NGBoost [16] to predict the parameters of each gene’s prior distribution from its features. Our gene features include gene expression levels, Gene Ontology terms, conservation across species, neural network embeddings of protein sequences, gene regulatory features, co-expression and protein-protein interaction features, sub-cellular localization, and intolerance to missense mutations (see Methods and Supplementary Note C for a full list).

Second, we use a model from population genetics to relate *s*_het_ to the observed LOF data (Figure 1**C**, right). This model allows us to fit the gradient-boosted trees for the prior by maximizing the likelihood of the LOF data. Specifically, we use the discrete-time Wright Fisher model with genic selection, a standard model in population genetics that accounts for mutation and genetic drift [13,17]. In our model, *s*_het_ is the reduction in fitness per copy of a LOF, and we infer *s*_het_ while keeping the mutation rates and demography fixed to values taken from the literature (Supplementary Note B). In particular, we assume that the average number of offspring an individual has is proportional to 1, 1 *− s*_het_, or 1 *−* 2*s*_het_ if they carry zero, one, or two copies of the LOF respectively, with these fitnesses lower bounded at zero. As such, if *s*_het_ is large, then individuals carrying a LOF allele will, on average, have fewer offspring either due to reduced viability or reduced fertility. Likelihoods are computed using new methods described in a companion paper [15].

Previous methods use either the number of *unique* LOFs or the sum of the frequencies of all LOFs in a gene, but we model the frequency of each individual LOF variant. We used LOF frequencies from the gnomAD consortium (v2), which consists of exome sequences from ~125,000 individuals for 19,071 protein-coding genes.

Combining these two components—the learned priors and the likelihood of the LOF data— we obtained posterior distributions over *s*_het_ for every gene. Throughout, we use the posterior mean value of *s*_het_ for each gene as a point estimate. While *s*_het_ is a quantitative measure of constraint, in Section 2.5 we provide qualitative descriptions of different ranges of *s*_het_ to aid practitioners in interpreting *s*_het_. See Methods for more details and Supplementary Table 2 for estimates of *s*_het_.

### 2.2 Population genetics model and gene features both affect the estimation of *s*_het_

First, we explored how LOF frequency and mutation rate relate to *s*_het_ in our population genetics model (Figure 2**A**). Invariant sites with high mutation rates are indicative of strong selection (*s*_het_ *>* 10^*−*2^), consistent with [18], while invariant sites with low mutation rates are consistent with essentially any value of *s*_het_ for the demographic model considered here. Regardless of mutation rate, singletons are consistent with most values of *s*_het_ but can rule out extremely strong selection, and variants observed at a frequency of *>*10% rule out even moderately strong selection (*s*_het_ *>* 10^*−*3^).

**Figure 2:**
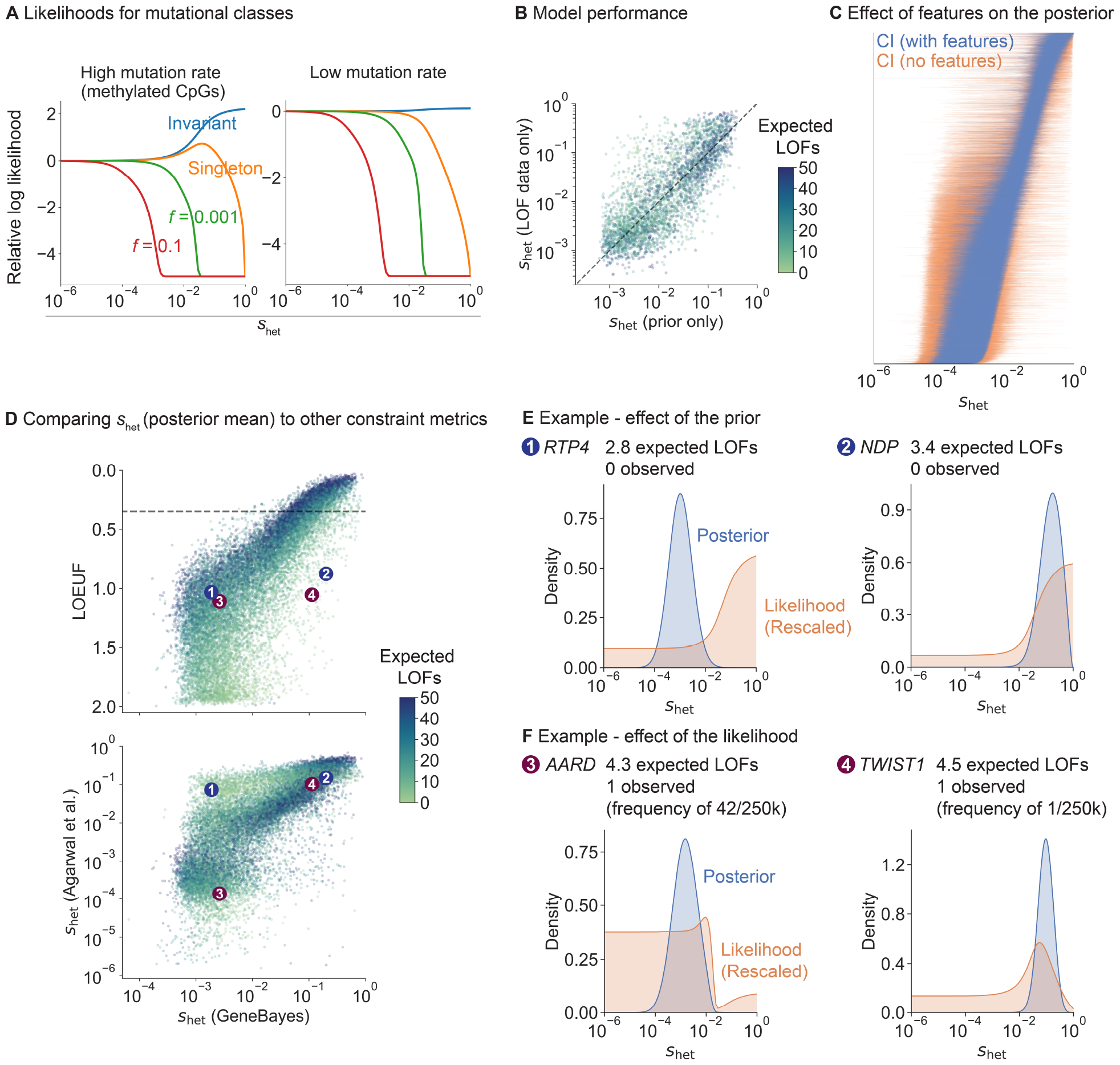
Factors that contribute to our estimates of s_het_. **A)** Likelihood curves for different allele frequencies (f) and mutation rates. **B)** Scatterplot of s_het_ estimated from LOF data (y-axis; posterior mean from a model without features) against the prior’s predictions of s_het_ (x-axis; mean of learned prior). Dotted line denotes y = x. Each point is a gene, colored by the expected number of LOFs. **C)** Comparison of posterior distributions of s_het_ (95% Credible Intervals) from a model with (blue lines) and without (orange lines) gene features. Genes are ordered by their posterior mean in the model with gene features. **D)** Top: scatterplot of LOEUF (y-axis) and our s_het_ estimates (x-axis; posterior mean). Each point is a gene, colored by the expected number of LOFs. Bottom: scatterplot of s_het_ estimates from [4] (y-axis; posterior mode) and our s_het_ estimates (x-axis; posterior mean). Numbered points refer to genes in panels **E** and **F. E)** RTP4 and NDP are two example genes where the gene features substantially affect the posterior. We plot their posterior distributions (blue) and likelihoods (orange; rescaled so that the area under the curve = 1). **F)** AARD and TWIST1 are two example genes with the same LOEUF but different s_het_. Posteriors and likelihoods are plotted as in panel **E**.

To assess how informative gene features are about *s*_het_, we trained our model on a subset of genes and evaluated the model on held-out genes (Figure 2**B**, Methods). We computed the Spearman correlation between *s*_het_ estimates from the prior and *s*_het_ estimated from the LOF data only. The correlation is high and comparable between train and test sets (Spearman *ρ* = 0.80 and 0.77 respectively), indicating the gene features alone are highly predictive of *s*_het_ and that this is not a consequence of overfitting.

To further characterize the impact of features on our estimates of *s*_het_, we removed all features from our model and recalculated posterior distributions (Figure 2**C**). For most genes, posteriors are substantially more concentrated when using gene features.

Some of our features are evolutionary measures of constraint, such as conservation among mammals, or the degree of constraint estimated from missense variants [19]. Given that these features may be correlated with LOF variation in a way independent of selection (e.g., local variation in mutation rate that is not well-captured by trinucleotide context), we wanted to make sure that these features were not majorly biasing our results. As such, we trained a version of our model that excluded these features, finding the results to be extremely concordant (Supplementary Figure 8**A**, Supplementary Note D).

We also made sure that our results were insensitive to the genetic ancestries of the individuals used when computing LOF frequencies by retraining our model using different subsets of the data (Supplementary Figure 6, Supplementary Note B).

Next, we compared our estimates of *s*_het_ using GeneBayes to LOEUF and to selection coefficients estimated by [4] (Figure 2**D**). To facilitate comparison, we use the posterior modes of *s*_het_ reported in [4] as point estimates, but we note that [4] emphasizes the value of using full posterior distributions. While the correlation between our estimates is high for genes with sufficient LOFs (for genes with more LOFs than the median, Spearman *ρ* with LOEUF = 0.94; *ρ* with *s*_het_ from [4] = 0.87), it is lower for genes with few expected LOFs (for genes with fewer LOFs than the median, Spearman *ρ* with LOEUF = 0.71; *ρ* with *s*_het_ from [4] = 0.69).

We further explored the reduced correlations for genes with few expected LOFs. For example, *RTP4* and *NDP* have few expected LOFs, and their likelihoods are consistent with any level of constraint (Figure 2**E**). Due to the high degree of uncertainty, LOEUF considers both genes to be unconstrained, while the *s*_het_ point estimates from [4] err in the other direction and consider both genes to be constrained (Figure 2**D**). This uncertainty arises from use of the LOF data alone, and is captured by the wide posterior distributions for the *s*_het_ estimates from [4]. In contrast, by using gene features, our posterior distributions of *s*_het_ indicate that *NDP* is strongly constrained but *RTP4* is not, consistent with the observation that hemizygous LOFs in *NDP* cause Norrie Disease, where degeneration of the neuroretina causes early childhood blindness [20].

In contrast to estimates of *s*_het_, LOEUF further ignores information about allele frequencies by considering only the number of unique LOFs, resulting in a loss of information. For example, *AARD* and *TWIST1* have almost the same numbers of observed and expected unique LOFs, so LOEUF is similar for both (LOEUF = 1.1 and 1.06 respectively). However, while *TWIST1*’s observed LOF is present in only 1 of 246,192 alleles, *AARD*’s is ~40*×* more frequent. Consequently, the likelihood rules out the possibility of strong constraint at *AARD* (Figure 2**F**), causing the two genes to differ in their estimated selection coefficients (Figure 2**D**).

In contrast, *TWIST1* has a posterior mean *s*_het_ of 0.11 when using gene features, indicating very strong selection. Consistent with this, TWIST1 is a transcription factor critical for specification of the cranial mesoderm, and heterozygous LOFs in the gene are associated with Saethre-Chotzen syndrome, a disorder characterized by congenital skull and limb abnormalities [21, 22].

As expected, genes with higher numbers of expected LOFs generally have greater concordance between their likelihoods and posterior distributions. We provide additional examples of genes with varying numbers of expected LOFs in Supplementary Figure 1.

Besides *NDP* and *TWIST1*, many genes are considered constrained by *s*_het_ but not by LOEUF, which is designed to be highly conservative. In Table 1, we list 15 examples in the top ~15% most constrained genes by *s*_het_ but in the ~75% least constrained genes by LOEUF, selected based on their clinical significance and prominence in the literature (Methods). One notable example is a set of 18 ribosomal protein genes for which heterozygous disruption causes Diamond-Blackfan anemia—a rare genetic disorder characterized by an inability to produce red blood cells [23] (Supplementary Table 1). Sixteen of the genes are considered strongly constrained by *s*_het_. In contrast, only 6 are considered constrained by LOEUF (LOEUF *<* 0.35), as many of these genes have few expected unique LOFs. Yet, collectively, these 18 proteins have ~139 expected unique LOFs but only 3 observed. If a single gene had this combination of observed and expected unique LOFs, it would have a LOEUF score of 0.06, consistent with extreme selective constraint. This highlights that LOEUF conflates lack of statistical power with a presumed lack of constraint.

**Table 1:** OMIM genes constrained by s_het_ but not by LOEUF. Mutations that disrupt the functions of these genes are associated with Mendelian diseases in the OMIM database [36]. Genes are ordered by s_het_ (posterior mean). Obs. and Exp. are the unique number of observed and expected LOFs respectively. *RPS15A is associated with Diamond-Blackfan anemia along with 12 other genes considered constrained by s_het_ but not by LOEUF (Supplementary Table 1), with 9 of the 12 genes falling outside the most constrained quartile by LOEUF. These genes were chosen from 301 genes that had s_het_ > 0.1 but were not in the most constrained LOEUF quartile. This includes 71 of 3,045 genes with pathogenic ClinVar variants that fall outside the most constrained LOEUF quartile.

| Gene | $s_{het}$ | LOEUF | Obs. | Exp. | Condition and reference |
| --- | --- | --- | --- | --- | --- |
| *RPS15A\** | 0.68 | 0.56 | 0 | 5.4 | *Diamond-Blackfan anemia*: Red blood cell aplasia resulting in growth, craniofacial, and other congenital defects [23] |
| *DCX* | 0.28 | 0.62 | 3 | 12.6 | *Lissencephaly*: Migrational arrest of neurons resulting in mental retardation and seizures [24] |
| *UBE2A* | 0.28 | 0.54 | 0 | 5.6 | *Intellectual disorder, Nascimento type*: Intellectual disability characterized by dysmorphic features [25] |
| *PQBP1* | 0.28 | 0.50 | 1 | 9.5 | *Renpenning syndrome*: Mental retardation with short stature and a small head size [26] |
| *NAA10* | 0.28 | 0.52 | 1 | 9.1 | *Syndromic microphthalmia*: Missing or abnormally small eyes from birth [27] |
| *SOX3* | 0.22 | 0.86 | 1 | 5.5 | *Intellectual disorder and isolated growth hormone deficiency*: Impaired fetal growth and intellectual development [28] |
| *NDP* | 0.20 | 0.88 | 0 | 3.4 | *Norrie disease*: Retinal dystrophy resulting in early childhood blindness, mental disorders, and deafness [20] |
| *EIF5A* | 0.19 | 0.54 | 1 | 8.7 | *Faundes-Banka syndrome*: Developmental delay, microcephaly, and facial dysmorphisms [29] |
| *CDKN1C* | 0.19 | 0.53 | 0 | 5.7 | *Beckwith-Wiedemann syndrome*: Pediatric overgrowth with predisposition to tumor development [30] |
| *BCAP31* | 0.15 | 0.65 | 2 | 9.7 | *Deafness, dystonia, and cerebral hypomyelination* Motor and intellectual disabilities, with deafness and involuntary muscle contraction [31] |
| *SOX2* | 0.14 | 0.57 | 1 | 8.3 | *Syndromic microphthalmia*: Missing or abnormally small eyes from birth [32] |
| *SH2D1A* | 0.14 | 0.96 | 1 | 4.9 | *Lymphoproliferative syndrome*: Immunodeficiency characterized by severe immune dysregulation after viral infection [33] |
| *GATA4* | 0.12 | 0.53 | 3 | 14.7 | *Atrial septal defect*: Congenital heart defect resulting in a hole between the atria [34] |
| *TWIST1* | 0.11 | 1.1 | 1 | 4.5 | *Saethre-Chotzen syndrome*: Craniosynostosis, facial dysmorphism, and hand and foot abnormalities [21] [22] |
| *TAFAZZIN* | 0.11 | 0.49 | 2 | 13.0 | *Barth syndrome*: Disorder in lipid metabolism characterized by heart, muscle, immune, and growth defects [35] |

### 2.3 Utility of s_het_ in prioritizing phenotypically important genes

To assess the accuracy of our *s*_het_ estimates and evaluate their ability to prioritize genes, we first used these estimates to classify genes essential for survival of human cells *in vitro*. Genome-wide CRISPR growth screens have measured the effects of gene knockouts on cell survival or proliferation, quantifying the *in vitro* importance of each gene for fitness [37,38]. We find that our estimates of *s*_het_ outperform other constraint metrics at classifying essential genes (Figure 3**A**, left; bootstrap *p <* 7 *×* 10^*−*7^ for pairwise differences in AUPRC between our estimates and other metrics). The difference is largest for genes with few expected LOFs, where *s*_het_ (GeneBayes) retains similar precision and recall while other metrics lose performance (Figure 3**A**, right). Our performance gains remain even when comparing to LOEUF computed using gnomAD v4, which contains roughly 6*×* as many individuals (Supplementary Figure 7**A**), highlighting that sharing information across genes is more important than increasing sample sizes, a point we made in [15]. In addition, our estimates of *s*_het_ outperform other metrics at classifying nonessential genes (Supplementary Figure 7**B**).

**Figure 3:**
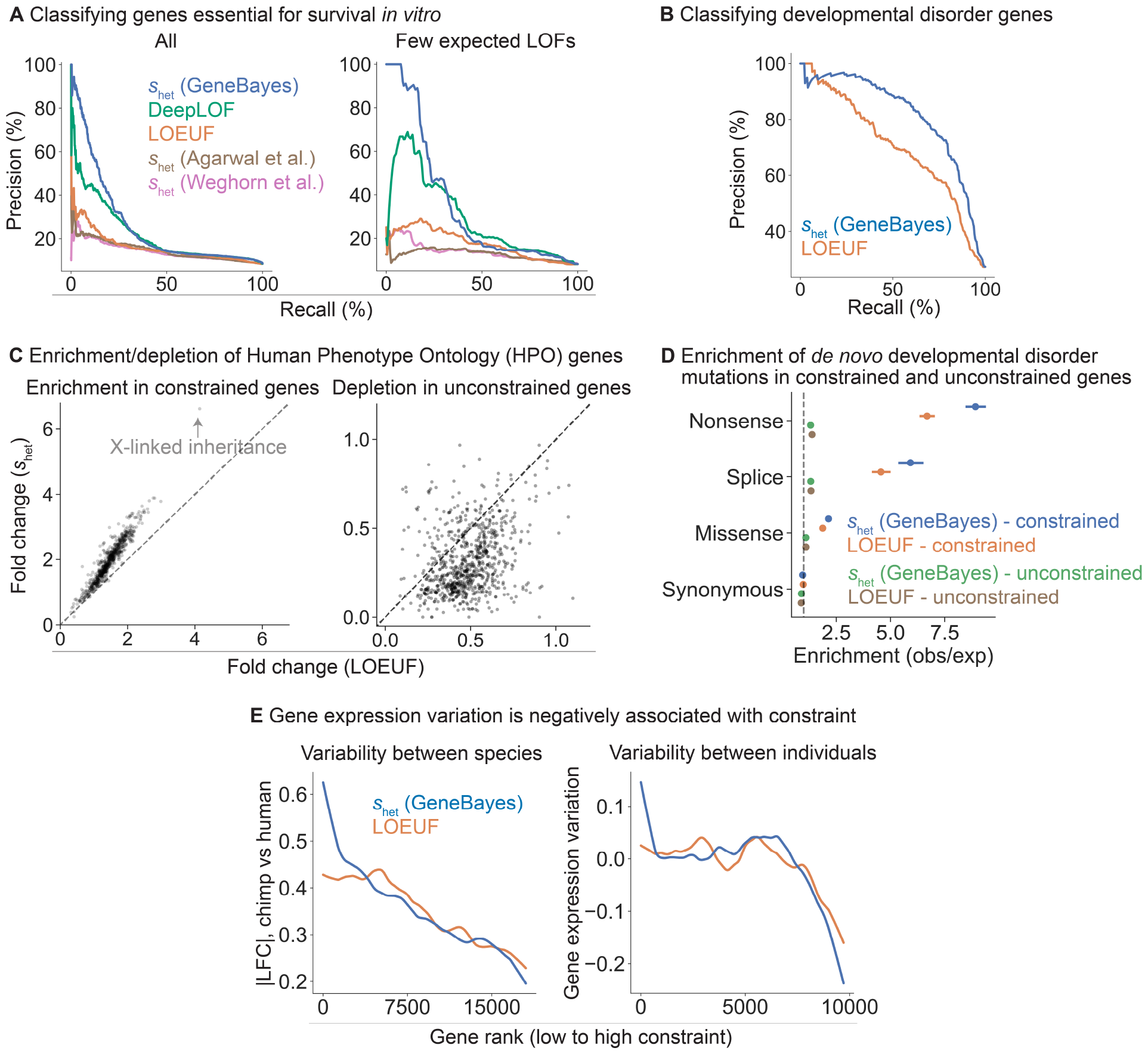
GeneBayes estimates of s_het_ perform well at identifying constrained and unconstrained genes. **A)** Precision-recall curves comparing the performance of s_het_ against other methods in classifying essential genes (left: all genes, right: quartile of genes with the fewest expected unique LOFs). **B)** Precision-recall curves comparing the performance of s_het_ against LOEUF in classifying developmental disorder genes. **C)** Scatterplots showing the enrichment (respectively depletion) of the top 10% most (respectively least) constrained genes in HPO terms, with genes ranked by s_het_ (y-axis) or LOEUF (x-axis). **D)** Enrichment of de novo mutations in patients with developmental disorders, calculated as the observed number of mutations over the expected number under a null mutational model. We plot the enrichment of synonymous, missense, splice, and nonsense variants in the 10% most constrained genes, ranked by s_het_ (blue) or LOEUF (orange); or enrichment in the remaining genes, ranked by s_het_ (green) or LOEUF (brown). Bars represent 95% confidence intervals. **E)** Left: LOESS curve showing the relationship between constraint (gene rank, x-axis) and absolute log fold change in expression between chimp and human cortical cells (y-axis). Genes are ranked by s_het_ (blue) or LOEUF (orange). Right: LOESS curve showing the relationship between constraint (gene rank, x-axis) and gene expression variation in GTEx samples after controlling for mean expression levels.

DeepLOF [14], the only other method that combines information from both LOF data and gene features, outperforms methods that rely exclusively on LOF data, highlighting the importance of using auxiliary information. Yet, DeepLOF uses only the number of unique LOFs, discarding frequency information. As a result, it is outperformed by our method, indicating that careful modeling of LOF frequencies also contributes to the performance of our approach.

Next, we performed further comparisons of our estimates of *s*_het_ against LOEUF, as LOEUF and its predecessor pLI are extremely popular metrics of constraint. To evaluate the ability of these methods to prioritize disease genes, we first used *s*_het_ and LOEUF to classify curated developmental disorder genes [39]. Here, *s*_het_ outperforms LOEUF (Figure 3**B**; bootstrap *p* = 5 *×* 10^*−*20^ for the difference in AUPRC) and performs favorably compared to additional constraint metrics (Supplementary Figure 7**C**).

We find that our estimates of *s*_het_ are not strongly dependent on any individually important features (Supplementary Figure 8**B,C**). In addition, *s*_het_ outperforms LOEUF even for genes with sufficient numbers of expected LOFs, although the measures become more concordant (Supplementary Figure 9).

Next, we considered a broader range of phenotypic abnormalities annotated in the Human Phenotype Ontology (HPO) [40]. For each HPO term, we calculated the enrichment of the 10% most constrained genes and depletion of the 10% least constrained genes, ranked using *s*_het_ or LOEUF. Genes considered constrained by *s*_het_ are 2.0-fold enriched in HPO terms, compared to 1.4-fold enrichment for genes considered constrained by LOEUF (Figure 3**C**, left). Additionally, genes considered unconstrained by *s*_het_ are 3.2-fold depleted in HPO terms, compared to 2.1-fold depletion for genes considered constrained by LOEUF (Figure 3**C**, right).

X-linked inheritance is one of the terms with the largest enrichment of constrained genes (6.7-fold enrichment for *s*_het_ and 4.1-fold enrichment for LOEUF). The ability of *s*_het_ to prioritize X-linked genes may prove particularly useful, as many disorders are enriched for X-chromosome genes [41] and the selection on losing a single copy of such genes is stronger on average [4]. Yet, population-scale sequencing alone has less power to detect a given level of constraint on X-chromosome genes, as the number of X chromosomes in a cohort with males is smaller than the number of autosomes.

We next assessed if *de novo* disease-associated variants are enriched in constrained genes, similar to the analyses in [4,5]. To this end, we used data from 31,058 trios to calculate for each gene the enrichment of *de novo* synonymous, missense, and LOF mutations in offspring with DDs relative to unaffected parents [5]. We found that for missense and LOF variants, enrichment is higher for genes considered constrained by *s*_het_, with the highest enrichment observed for LOF variants (Figure 3**D**; enrichment of *s*_het_ and LOEUF respectively, for missense mutations = 2.1, 1.9; splice site mutations = 5.9, 4.6; and nonsense mutations = 8.9, 6.7). Synonymous variants are not enriched in genes constrained by either method. Consistent with previous findings, the excess burden of *de novo* variants is predominantly in highly constrained genes (Figure 3**D**). Notably, this difference in enrichment remains after removing known DD genes (Supplementary Figure 7**D**, right). Together, these results indicate that *s*_het_ not only improves identification of known disease genes but may also facilitate discovery of novel DD genes [5].

In addition to rare *de novo* disease-associated variants, we find that common variant heritability as computed using stratified LD score regression is enriched in constrained genes (Supplementary Figure 7**E**), consistent with the findings from [5]. For 380 of 438 highly-heritable traits (87%), heritability is more highly enriched in the decile of genes most highly constrained by *s*_het_ than the decile most highly constrained by LOEUF (Supplementary Figure 7**E**, Methods), with a mean enrichment across traits of 1.5-fold.

Finally, constraint can also be related to longer-term evolutionary processes that give rise to the variation among individuals or species, including variation in gene expression levels. We expect constrained genes to maintain expression levels closer to their optimal values across evolutionary time scales, as each LOF can be thought of as a ~50% reduction in expression. Consistent with this expectation, we find that less constrained genes have larger absolute differences in expression between human and chimpanzee in cortical cells [42], with a stronger correlation for *s*_het_ than for LOEUF (Figure 3**E**). This pattern should also hold when considering the variation in expression within a species. We quantified variance in gene expression levels estimated from RNA-seq samples in GTEx [43] after controlling for mean expression levels, and found that the variance decreases with increased constraint, again with a stronger correlation for *s*_het_ (Figure 3**E**; Methods).

### 2.4 Interpreting the learned relationship between gene features and s_het_

Our framework allows us to learn the relationship between gene features and *s*_het_ in a statistically principled way. In particular, by fitting a model with all of the features jointly, we can account for dependencies between the features. To interrogate the relationship between features and *s*_het_, we divided our gene features into 10 distinct categories (Figure 4**A**) and trained a separate model per category using only the features in that category. We found that missense constraint, gene expression patterns, evolutionary conservation, and protein embeddings are the most informative categories.

**Figure 4:**
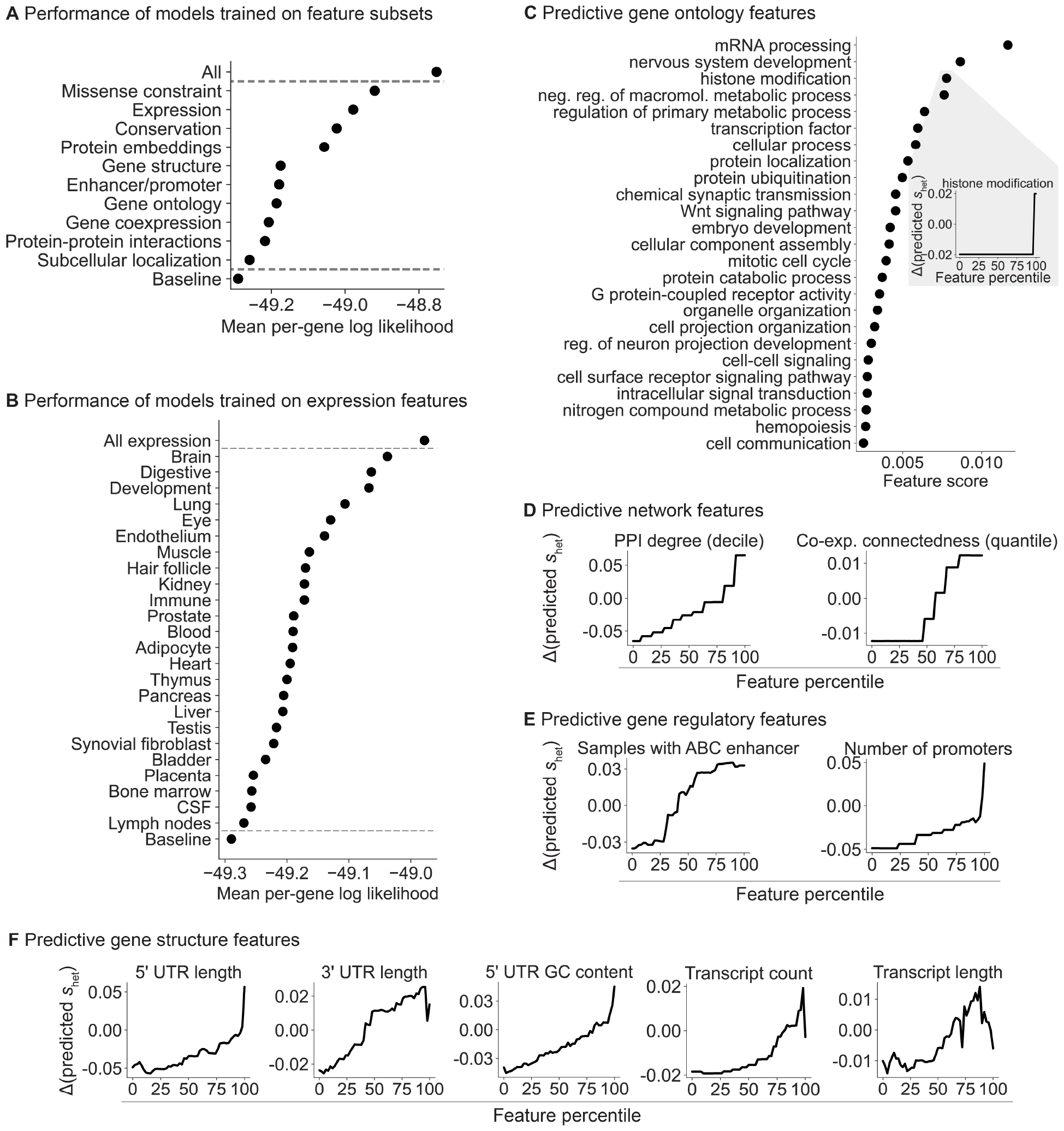
Breakdown of the gene features important for s_het_ prediction. **A)** Ordered from highest to lowest, plot of the mean per-gene log likelihood over the test genes for models separately trained on categories of features. “All” and “Baseline” include all and no features respectively. **B)** Plot of the mean per-gene log likelihood, as in panel **A**, for models separately trained on expression features grouped by tissue, cell type, or developmental stage. **C)** Ordered from highest to lowest, feature scores for individual gene ontology (GO) terms. Inset: lineplot showing the change in predicted s_het_ for a feature as the feature value is varied. **D)** Lineplot as in panel **C** (inset) for protein-protein interaction (PPI) and co-expression features, **E)** enhancer and promoter features, and **F)** gene structure features.

Next, we further divided the expression features into 24 subgroups, representing tissues, cell types, and developmental stage (Table 6). Expression patterns in the brain, digestive system, and during development are the most predictive of constraint (Figure 4**B**). Notably, a study that matched Mendelian disorders to tissues through literature review found that a sizable plurality affect the brain [44]. Meanwhile, most of the top digestive expression features are also related to development (e.g., expression component loadings in a fetal digestive dataset [45]). The importance of developmental features is consistent with the severity of many developmental disorders and the expectation that selection is stronger on early-onset phenotypes [46], supported by the findings of [4].

To quantify the relationship between constraint and individual features, we changed the value of one feature at a time and used the variation in predicted *s*_het_ over the feature values as the score for each feature (Methods).

We first explored some of the individual Gene Ontology (GO) terms most predictive of constraint (Figure 4**C**). Consistent with the top expression features, the top GO features highlight developmental and brain-specific processes as important for selection.

Next, we analyzed network (Figure 4**D**), gene regulatory (Figure 4**E**), and gene structure (Figure 4**F**) features. Protein-protein interaction (PPI) and gene co-expression networks have highlighted “hub” genes involved in numerous cellular processes [47,48], while genes linked to GWAS variants have more complex enhancer landscapes [49]. Consistent with these studies, we find that connectedness in PPI and co-expression networks as well as enhancer and promoter count are positively associated with constraint (Figure 4**D,E**). In addition, gene structure affects gene function—for example, UTR length and GC content affect RNA stability, translation, and localization [50, 51]—and likewise, several gene structure features are predictive of constraint (Figure 4**F**), consistent with recent work on UTRs [52]. Our results indicate that more complex genes—genes that are involved in more regulatory connections, that are more central to networks, and that have more complex gene structures—are generally more constrained.

Gene length is predictive of *s*_het_ (Figure 4**F**), but also correlates with the amount of information in the LOF data as well as a number of other gene features (Supplementary Figure 10**A,B,C**). While the model learns the importance of all features jointly, and hence could adjust for gene length when considering other features, we wanted to be sure that the signal from other features was not generally driven by their correlation with gene length. As such, we computed partial correlations between each feature and posterior mean *s*_het_ adjusting for gene length, and found that gene length explains at most a modest amount of the correlation between most features and *s*_het_ (Supplementary Figure 10**D**).

### 2.5 Contextualizing the strength of selection against gene loss-of-function

A major benefit of *s*_het_ over LOEUF and pLI is that *s*_het_ has a precise, intrinsic meaning in terms of fitness [1–4]. This facilitates comparison of *s*_het_ between genes, populations, species, and studies. For example, *s*_het_ can be compared to selection estimated from mutation accumulation or gene deletion experiments performed in model organisms [53,54]. More broadly, selection applies beyond LOFs. While we focused on estimating changes in fitness due to LOFs, consequences of non-coding, missense, and copy number variants can be understood through the same framework, as we expect such variants to also be under negative selection [18] due to ubiquitous stabilizing selection on traits [55]. Quantifying differences in the selection on variants will deepen our understanding of the evolution and genetics of human traits (see Discussion).

To contextualize our *s*_het_ estimates, we compared the distributions of *s*_het_ for different gene sets (Figure 5**A**) and genes (Figure 5**B**), and analyzed them in terms of selection regimes. To define such regimes, we first conceptualized selection on variants as a function of their effects on expression (Figure 5**C**), where heterozygous LOFs reduce expression by ~50% across all contexts relevant to selection. Under this framework, we can directly compare *s*_het_ to selection on other variant types— for the hypothetical genes in Figure 5**C**, a GWAS hit affecting Gene 1 has a stronger selective effect than a LOF affecting Gene 2, despite having a smaller effect on expression.

**Figure 5:**
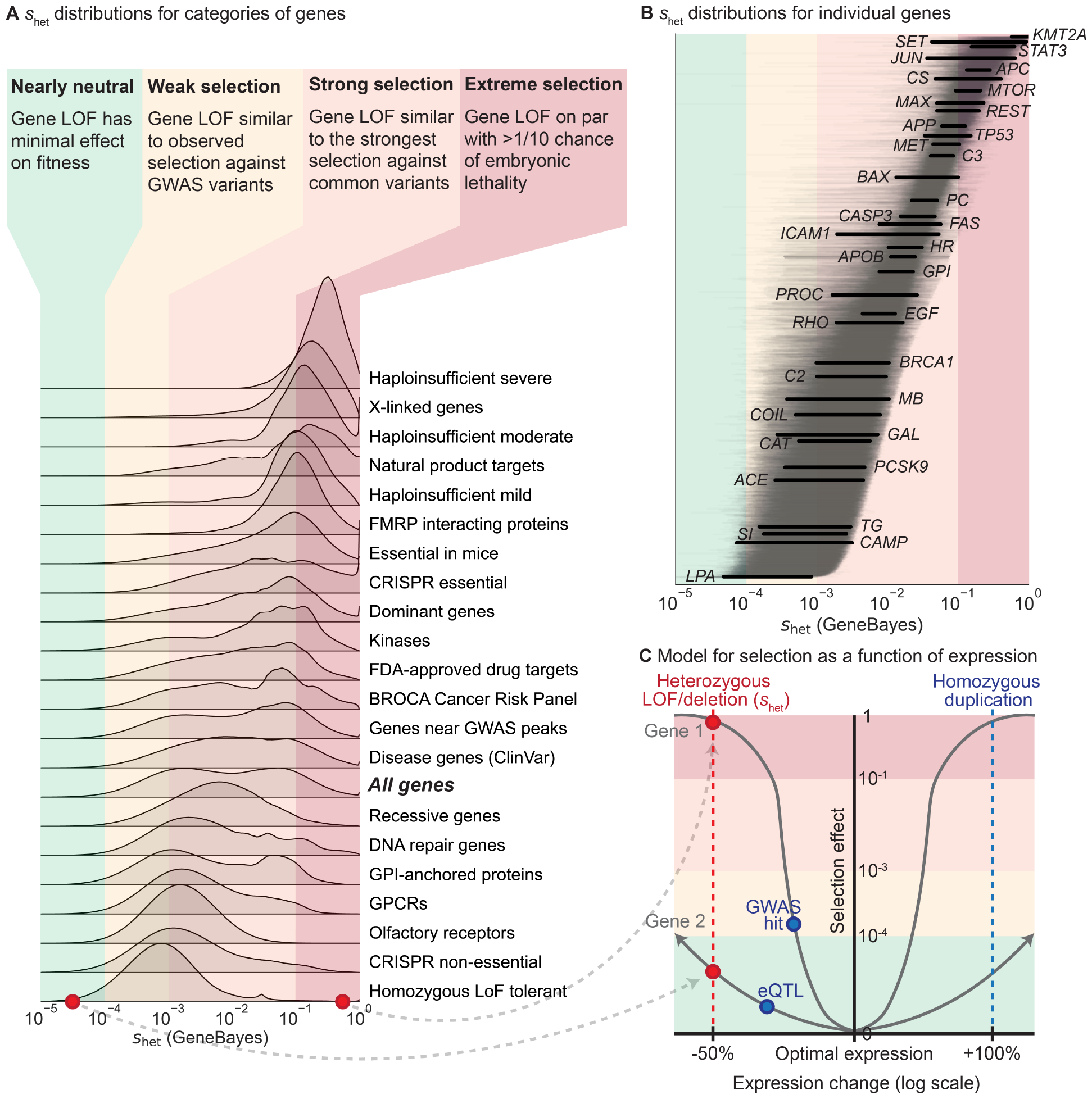
Comparing selection on LOFs (s_het_) between genes and to selection on other variant types. **A)** Distributions of s_het_ for gene sets, calculated by averaging the posterior distributions for the genes in each gene set. Gene sets are sorted by the mean of their distributions. Colors represent four general selection regimes. **B)** Posterior distributions of s_het_ for individual genes, ordered by mean. Lines represent 95% credible intervals, with labeled genes represented by thick black lines. Colors represent the selection regimes in panel **A. C)** Schematic demonstrating the hypothesized relationship between changes in expression (x-axis, log_2_ scale) and selection (y-axis) against these changes for two hypothetical genes, assuming stabilizing selection. The shapes of the curves are not estimated from real data. Background colors represent the selection regimes in panel **A**. The red points and line represent the effects of heterozygous LOFs and deletions on expression and selection, while the blue points and line represent the potential effects of other types of variants.

Next, we divided the range of possible *s*_het_ values into four regimes determined by theoretical considerations [56] and comparisons to other types of variants [57, 58]—nearly neutral, weak selection, strong selection, and extreme selection. LOFs in nearly neutral genes (*s*_het_ *<* 10^*−*4^) have minimal effects on fitness—the frequency of such variants is dominated by genetic drift rather than selection [56]. Under the weak selection regime (*s*_het_ from 10^*−*4^ to 10^*−*3^), gene LOFs have similar effects on fitness as typical GWAS hits, which usually have small or context-specific effects on gene expression or function [57]. Under the strong selection regime (*s*_het_ from 10^*−*3^ to 10^*−*1^), gene LOFs have fitness effects on par with the strongest selection coefficients measured for common variants, such as the selection estimated for adaptive mutations in *LCT* [58]. Finally, for genes in the extreme selection regime (*s*_het_ *>* 10^*−*1^), LOFs have an effect on fitness equivalent to a *>*2% chance of embryonic lethality, indicating that such LOFs have an extreme effect on survival or reproduction.

Gene sets vary widely in their constraint. For example, genes known to be haploinsufficient for severe diseases are almost all under extreme selection. In contrast, genes that can tolerate homozygous LOFs are generally under weak selection. One notable example of such a gene is *LPA*—while high expression levels are associated with cardiovascular disease, low levels have minimal phenotypic consequences [59, 60], consistent with limited conservation in the sequence or gene expression of *LPA* across species and populations [61, 62]

Other gene sets have much broader distributions of *s*_het_ values. For example, manually curated recessive genes are under weak to strong selection, indicating that many such genes are either not fully recessive or have pleiotropic effects on other traits under selection. For example, homozygous LOFs in *PROC* can cause life-threatening congenital blood clotting [63], yet *s*_het_ for *PROC* is non-negligible (Figure 5**B**), consistent with observations that heterozygous LOFs can also increase blood clotting and cause deep vein thrombosis [64].

Similarly, *s*_het_ values for ClinVar disease genes [65] span the range from weak to extreme selection, with only moderate enrichment for greater constraint relative to all genes. Consistent with this, the effects of disease on fitness depend on disease severity, age-of-onset, and prevalence throughout human history. For example, even though heterozygous loss of *BRCA1* greatly increases risk of breast and ovarian cancer [66], *BRCA1* is under strong rather than extreme selection. Possible partial explanations are that these cancers have an age-of-onset past reproductive age and are less prevalent in males, or that *BRCA1* is subject to some form of antagonistic pleiotropy [67, 68].

## 3 Discussion

Here, we developed an empirical Bayes approach to accurately infer *s*_het_, an interpretable metric of gene constraint. Our approach uses powerful machine learning methods to leverage vast amounts of functional and evolutionary information about each gene while coupling them to a population genetics model.

There are two advantages of this approach. First, the additional data sources result in substantially better performance than LOEUF across tasks, from classifying essential genes to identifying pathogenic *de novo* mutations. These improvements are especially pronounced for the large fraction of genes with few expected LOFs, where LOF data alone is underpowered for estimating constraint.

Second, by inferring *s*_het_, our estimates of constraint are interpretable in terms of fitness, and we can directly compare the impact of a loss-of-function across genes, populations, species, and studies.

As a selection coefficient, *s*_het_ can also be directly compared to other selection coefficients, even for different types of variants [3, 4]. In general, we believe genes are close to their optimal levels of expression and experience stabilizing selection [55], in which case expression-altering variants decrease fitness, with larger perturbations causing greater decreases (Figure 5**C**). Estimating the fitness consequences of other types of expression-altering variants, such as duplications or eQTLs, will allow us to map the relationship between genetic variation and fitness in detail, deepening our understanding of the interplay of expression, complex traits, and fitness [10, 57, 69, 70].

A recent method, DeepLOF [14], uses a similar empirical Bayes approach, but by estimating constraint from the number of observed and expected unique LOFs, it inherits the same difficulties regarding interpretation as pLI and LOEUF, and loses information by not considering variant frequencies. Another line of work [1, 2], culminating in [4], solved the issues with interpretability by directly estimating *s*_het_. Yet, by relying exclusively on LOFs, these estimates are underpowered for ~25% of genes. Furthermore, by using the aggregate frequencies of all LOF variants, previous *s*_het_ estimates [1, 2, 4] are not robust to misannotated LOF variants. Our approach eliminates this tradeoff between power and interpretability present in existing metrics.

Similar insights that combine evolutionary modeling and genomic features have been used to estimate constraint on non-coding variation [71–74], and extending our approach to non-coding variation would be an interesting direction for future work.

Our estimates of *s*_het_ will be useful for many applications. For example, by informing gene-level priors, LOEUF, pLI, and previous estimates of *s*_het_ have been used to increase the power of association studies based on rare or *de novo* mutations [5, 6, 75]. In such contexts, our *s*_het_ estimates can be used as a drop-in replacement. Additionally, extremely constrained and unconstrained genes may be interesting to study in their own right. Genes of unknown function with particularly high values of *s*_het_ should be prioritized for further study. Investigating highly constrained genes may give insights into the mechanisms by which cellular and organism-level phenotypes affect fitness [76].

While we primarily used the posterior means of *s*_het_ here, our approach provides the entire posterior distribution per gene, similar to [4]. In some applications, different aspects of the posterior may be more relevant than the mean. For example, when prioritizing rare variants for followup in a clinical setting, the posterior probability that *s*_het_ is high enough for the variant to severely reduce fitness may be more relevant.

As more exomes are sequenced, one might expect that we would be better able to more accurately estimate *s*_het_. Yet, in a companion paper [15], we show that increasing the sample size used for estimating LOF frequencies will provide essentially no additional information for the ~85% of genes with the lowest values of *s*_het_. This fundamental limit on how much we can learn about these genes from LOF data alone highlights the importance of approaches like ours that can leverage additional data types. By sharing information across genes, we can overcome this fundamental limit on how accurately we can estimate constraint.

Here we focused on estimating *s*_het_, but our empirical Bayes framework, GeneBayes, can be used in any setting where one has a model that ties a gene-level parameter to gene-level observable data (Supplementary Note E). For example, GeneBayes can be used to find trait-associated genes using variants from case/control studies [77, 78], or to improve power to find differentially expressed genes in RNA-seq experiments [79]. We provide a graphical overview of how GeneBayes can be applied more generally in Figure 6. Briefly, GeneBayes requires users to specify a likelihood model and the form of a prior distribution for their parameter of interest. Then, using empirical Bayes and a set of gene features, it improves power to estimate the parameter by flexibly sharing information across similar genes.

**Figure 6:**
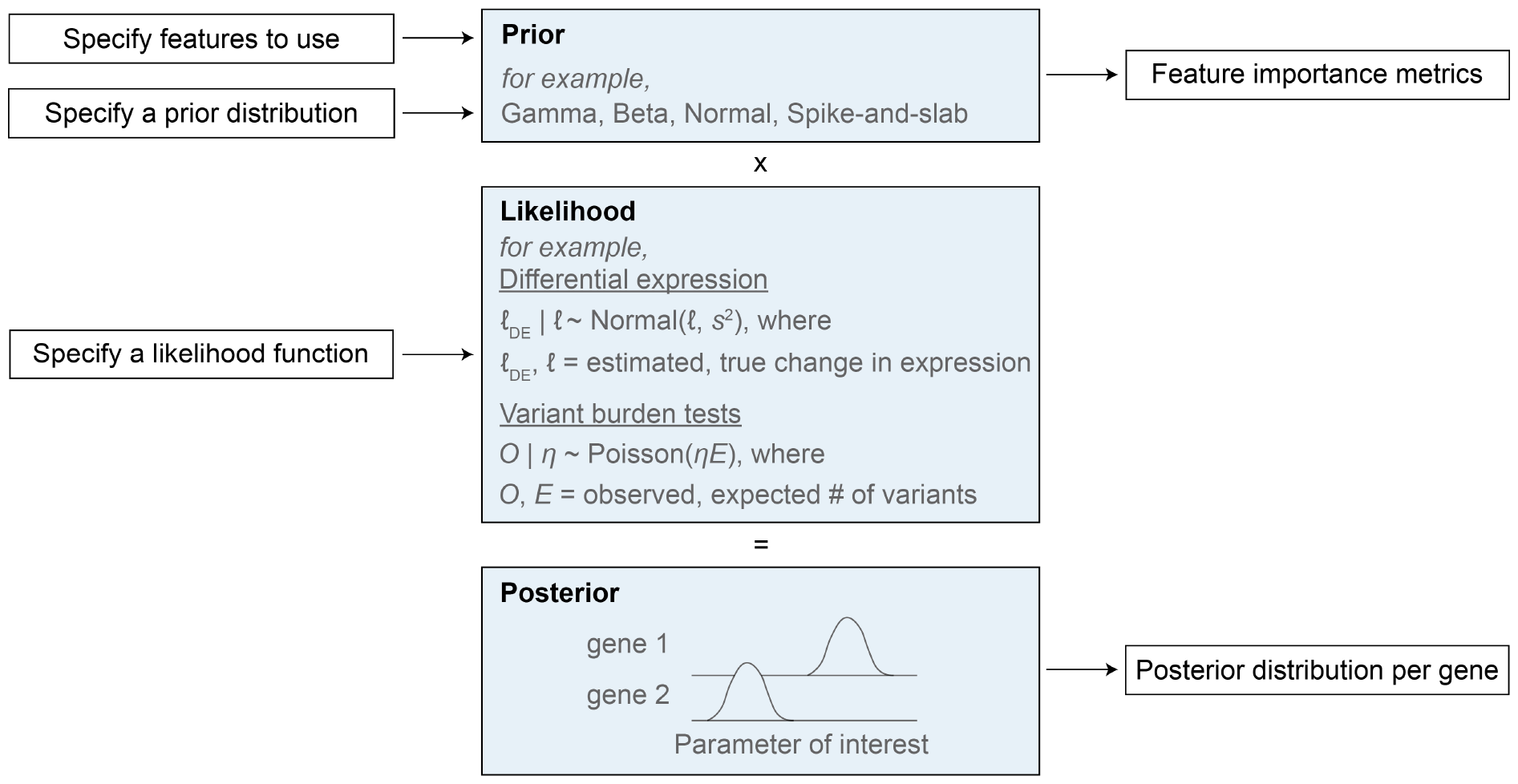
GeneBayes is a flexible framework for estimating gene-level properties. Schematic for how GeneBayes can be applied to estimate gene-level properties beyond s_het_, showing the key inputs and outputs and two example applications. See Supplementary Note E for more details.

In summary, we developed a powerful framework for estimating a broadly applicable and readily interpretable metric of constraint, *s*_het_. Our estimates provide a more informative ranking of gene importance than existing metrics, and our approach allows us to interrogate potential causes and consequences of natural selection.

## Data availability

Posterior means and 95% credible intervals for *s*_het_ are available in Supplementary Table 2. Posterior densities for *s*_het_ are available in Supplementary Table 3. A description of the gene features is available in Supplementary Table 4. These supplementary tables are also available at [80], along with likelihoods for *s*_het_, LOF variants with misannotation probabilities, and gene feature tables.

## Code availability

GeneBayes and code for estimating *s*_het_ are available at https://github.com/tkzeng/GeneBayes.

## Acknowledgements

We would like to thank Ipsita Agarwal, Molly Przeworski, Jesse Engreitz, and members of the Pritchard Lab for valuable feedback and discussions. This work was supported by NIH grants R01AG066490, R01HG011432, R01HG008140, and U01HG009431.

## 4 Methods

### Empirical Bayes overview

Many genes have few observed loss-of-function variants, making it challenging to infer constraint without additional information. Bayesian approaches that specify a prior distribution for each gene can provide such information to improve constraint estimates, but specifying prior distributions is challenging as we have limited prior knowledge about the selection coefficients, *s*_het_. Empirical Bayes procedures allow us to learn a prior distribution for each gene by combining information across genes.

To use the information contained in the gene features, we learn a mapping from a gene’s features to a prior specific for that gene. We parameterize this mapping using gradient-boosted trees, as implemented in `NGBoost` [16]. Intuitively, this approach learns a notion of “similarity” between genes based on their features, and then shares information across similar genes to learn how *s*_het_ relates to the gene features. This approach has two major benefits. First, by sharing information between similar genes, it can dramatically improve the accuracy of the predicted *s*_het_ values, particularly for genes with few expected LOFs. Second, by leveraging the LOF data, this approach allows us to learn about how the various gene features relate to fitness, which cannot be modeled from first principles.

For a more in-depth description of our approach along with mathematical and implementation details, see Supplementary Note A.

### Population genetic likelihood

To model how *s*_het_ relates to the frequency of individual LOF variants, we used the discrete-time Wright-Fisher model, with an approximation of diploid selection with additive fitness effects. We used a composite likelihood approach, assuming independence across individual LOF variants to obtain gene-level likelihoods. Within this composite likelihood, we model each individual variant as either having a selection coefficient of *s*_het_ with probability 1 *− p*_miss_, or having a selection coefficient of 0 with probability *p*_miss_. That is, *p*_miss_ acts as the prior probability that a given variant is misannotated, and we assume that misannotated variants evolve neutrally regardless of the strength of selection on the gene. All likelihoods were computed using new machinery developed in a companion paper [15].

Our model depends on a number of parameters—a demographic model of past population sizes, mutation rates for each site, and the probability of misannotation. The demographic model is taken from the literature [81] with modifications as described in [4]. The mutation rates account for trinucleotide context as well as methylation status at CpGs [12]. Finally, we estimated the probability of misannotation from the data.

For additional technical details and intuition see Supplementary Note B.

### Curation of LOF variants

We obtained annotations for the consequences of all possible single nucleotide changes to the hg19 reference genome from [82]. The effects of variants on protein function were predicted using Variant Effect Predictor (VEP) version 85 [83] using GENCODE v19 gene annotations [84] as a reference. We defined a variant as a LOF if it was predicted by VEP to be a splice acceptor, splice donor, or stop gain variant. In addition, predicted LOFs were further annotated using LOFTEE [12], which implements a series of filters to identify variants that may be misannotated (for example, LOFTEE considers predicted LOFs near the ends of transcripts as likely misannotations). For our analyses, we only kept predicted LOFs labelled as High Confidence by LOFTEE, which are LOFs that passed all of LOFTEE’s filters.

Next, we considered potential criteria for further filtering LOFs: cutoffs for the median exome sequencing read depth, cutoffs for the mean pext (proportion expressed across transcripts) score [82], whether to exclude variants that fall in segmental duplications or regions with low mappability [85], and whether to exclude variants flagged by LOFTEE as potentially problematic but that passed LOFTEE’s primary filters.

We trained models with these filters one at a time and in combination, and chose the model that had the best AUPRC in classifying essential from nonessential genes in mice. The filters we evaluated and chose for the final model are reported in Table 2. Since we used mouse gene essentiality data to choose the filters, we do not further evaluate *s*_het_ on these data.

**Table 2:** Filtering criteria for LOF curation.

| Filtering criterion | Tested values | Best value |
| --- | --- | --- |
| Cutoff for sequencing read depth (median across exomes) | 0×, 5×, 10×, 20× | 0× |
| Cutoff for mean pext across tissues | 0.05, 0.1 | 0.05 |
| Filter if variant falls in a segmental duplication or low mappability region | True, False | True |
| Filter if variant is flagged as potentially problematic | True, False | True |

**Table 3:**
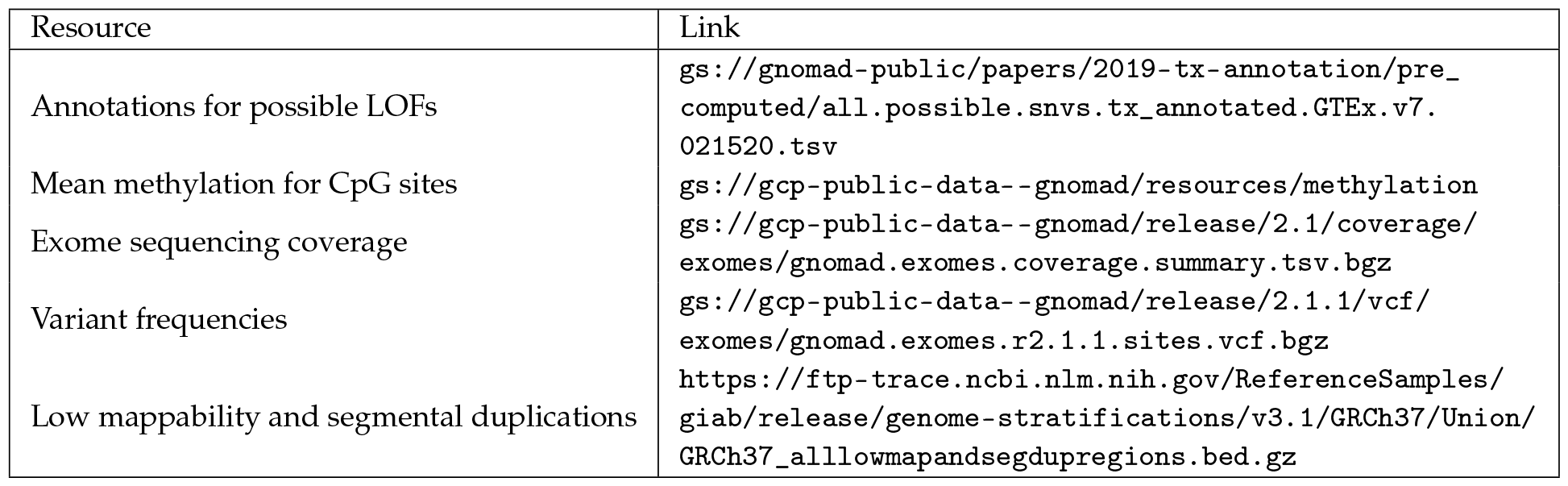
Sources for LOF data.

**Table 4:** Parameters for fitting the gradient-boosted trees.

| Parameter(s) | Tested values | Best value |
| --- | --- | --- |
| Learning rate | $2.5 \times 10^{-3}$, 0.01, 0.04 | 0.04 |
| Maximum tree depth (max_depth) | 3, 4, 5 | 3 |
| Data subsampling ratio (subsample) | 0.6, 0.8, 1 | 0.8 |
| Minimum weight of a leaf node (min_child_weight) | 1, 2, 4 | 4 |
| L1 regularization (alpha) | 1, 2, 4 | 2 |
| L2 regularization (lambda) | 0, 1, 2 | 0 |
| Number of trees to fit per iteration (n_estimators) | 1, 2, 4 | 1 |

We considered genes to be essential in mice if they are heterozygous lethal, as determined by [12] using data from heterozygous knockouts reported in Mouse Genome Informatics [86]. We classify genes as nonessential if they are reported as `Homozygous-Viable` or `Hemizygous-Viable` by the International Mouse Phenotyping Consortium [87] (annotations downloaded on 12/08/22 from https://www.ebi.ac.uk/mi/impc/essential-genes-search/).

Finally, we annotated each variant with its frequency in the gnomAD v2.1.1 exomes [12], a dataset of 125,748 uniformly-analyzed exomes that were largely curated from case–control studies of common adult-onset diseases. gnomAD provides precomputed allele frequencies for all variants that they call.

For potential LOFs that are not segregating, gnomAD does not release the number of individuals that were genotyped at those positions. For these sites, we used the median number of genotyped individuals at the positions for which gnomAD does provide this information. We performed this separately on the autosomes and X chromosome.

Data sources for the variant annotations, filters, and frequencies, as well as additional information used to compute likelihoods are listed in Table 3.

### Feature processing and selection

We compiled 10 types of gene features from several sources:

1. Gene structure (e.g., number of transcripts, number of exons, GC content)
2. Gene expression across tissues and cell lines
3. Biological pathways and Gene Ontology terms
4. Protein-protein interaction networks
5. Co-expression networks
6. Gene regulatory landscape (e.g., number and properties of enhancers and promoters)
7. Conservation across species
8. Protein embeddings
9. Subcellular localization
10. Missense constraint

Additionally, we included an indicator variable that is 1 if the gene is on the non-pseudoautosomal region of the X chromosome and 0 otherwise.

For a description of the features within each category and where we acquired them, see Supplementary Note C.

### Training and validation

We fine-tuned a set of hyperparameters for our full empirical Bayes approach, using the best hyperparameters from an initial feature selection step (described in Supplementary Note C) as a starting point. To minimize overfitting, we split the genes into three sets—a training set (chromosomes 7-22, X), a validation set for hyperparameter tuning (chromosomes 2, 4, 6), and a test set to evaluate overfitting (chromosomes 1, 3, 5). During each training iteration, one or more trees were added to the model to fit the gradient of the loss on the training set. We stopped model training once the loss on the validation set did not improve for 10 iterations in a row (or the maximum number of iterations, 1,000, was reached). Using this approach, we performed a grid search over the hyperparameters listed in Table 4, and used the combination with the lowest validation loss and best performance at classifying mouse essential genes (mean of the ranks on the two metrics).

### Choosing genes for Table 1

To identify genes that are considered constrained by *s*_het_ but not by LOEUF, we filtered for genes with *s*_het_ *>* 0.1 (top ~15% most constrained genes, analogous to the recommended LOEUF cutoff of 0.35 [67], which corresponds to the top ~16% of genes) and LOEUF *>* 0.47 (least constrained ~75% of genes). Of these, we identified genes where heterozygous or hemizygous mutations that decrease the amount of functional protein (e.g. LOF mutations) are associated with Mendelian disorders in the Online Mendelian Inheritance in Man (OMIM) database [36]. We chose genes for Table 1 primarily based on their prominence in the existing literature.

We define a gene as having a pathogenic variant in ClinVar if it contains a variant annotated with `CLNSIG` = `Pathogenic`. We downloaded ClinVar variants from https://ftp.ncbi.nlm.nih.gov/pub/clinvar/vcf_GRCh38/ on 12/03/2023.

## Evaluation on additional datasets

### Definition of human essential and nonessential genes

We obtained data from 1,085 CRISPR knockout screens quantifying the effects of genes on cell survival or proliferation from the DepMap portal (22Q2 release) [37, 38]. Scores from each screen are normalized such that nonessential genes identified by [88] have a median score of 0 and that common essential genes identified by [88, 89] have a median score of *−*1.

In classifying essential genes (Figure 3**A**), we define a gene as essential if its score is *< −* 1 in at least 25% of screens, and as *not* essential if its score is *> −* 1 in all screens. In classifying nonessential genes, we define a gene as nonessential if it has a minimal effect on growth in most cell lines (absolute effect *<*0.25 in at least 99% of screens), and as *not* nonessential if its score is *<*0 in all screens.

### Definition of developmental disorder genes

Through the Deciphering Developmental Disorders (DDD) study [39], clinicians have annotated a subset of genes with the strength and nature of their association with developmental disorders. We classify genes as developmental disorder genes if they are annotated by the DDD study with `confidence_category` = `definitive` and `allelic_requirement` = `monoallelic_autosomal`, `monoallelic_X_hem` (hemizygous), or `monoallelic_X_het` (heterozygous).

We classify genes as not associated with developmental disorders if they are annotated by the DDD study, do not meet the above criteria for association with a disorder, and are not annotated with `confidence_category` = `strong`, `moderate`, or `limited` and `allelic_requirement` = `monoallelic_autosomal`, `monoallelic_X_hem`, or `monoallelic_X_het`.

We downloaded genes with DDD annotations from https://www.deciphergenomics.org/ddd/ddgenes on 11/19/2023.

### Enrichment/depletion of Human Phenotype Ontology (HPO) genes

The Human Phenotype Ontology (HPO) provides a structured organization of phenotypic abnormalities and the genes associated with them, with each HPO term corresponding to a phenotypic abnormality. We calculated the enrichment of constrained genes in each HPO term with at least 200 genes as the ratio (fraction of HPO genes under constraint)/(fraction of background genes under constraint). We defined genes under constraint to be the decile of genes considered most constrained by *s*_het_ or LOEUF. To choose background genes, we sampled from the set of all genes to match each HPO term’s distribution of expected unique LOFs. Similarly, we calculated the depletion of unconstrained genes in each HPO term as the ratio (fraction of HPO genes not under constraint)/(fraction of background genes not under constraint), where we define genes not under constraint to be the decile of genes considered least constrained by *s*_het_ or LOEUF.

We downloaded HPO phenotype-to-gene annotations from http://purl.obolibrary.org/obo/hp/hpoa/phenotype_to_genes.txt on 01/27/2023.

### Enrichment of *de novo* mutations in developmental disorder patients

We used the enrichment metric developed by [5] in their analysis of *de novo* mutations (DNMs) identified from exome sequencing of 31,058 developmental disorder patients and their unaffected parents. Enrichment of DNMs in developmental disorder patients was calculated as the ratio of observed DNMs in patients over the expected number under a null mutational model that accounts for the study sample size and triplet mutation rate at the mutation sites [90].

For Figure 3**D**, we calculated the enrichment of DNMs in constrained genes, defined as the decile of genes considered most constrained by *s*_het_ or LOEUF. For Supplementary Figure 7**D**, we calculated the enrichment of DNMs in constrained genes with and without known associations with development disorders. We defined a gene as having a known association if it is annotated by the DDD study (see Methods section “Definition of developmental disorder genes”) with `confidence_category` = `definitive` or `strong` and `allelic_requirement` = `monoallelic_autosomal`, `monoallelic_X_hem` (hemizygous), or `monoallelic_X_het` (heterozygous).

For each set of genes, we computed the mean enrichment over sites and 95% Poisson confidence intervals for the mean using the code provided by [5].

### Heritability enrichment in constrained genes

We computed the heritability enrichment in the top 10% of genes constrained by *s*_het_ or LOEUF using stratified LD score regression (S-LDSC) [91]. To do this, we divided the heritability enrichment in constrained genes as reported by S-LDSC by the heritability enrichment in all genes. We linked variants to genes if they were in or within 100kb of the gene body, and ran S-LDSC using 1000G EUR Phase3 genotype data to estimate LD scores, baseline v2.2 annotations, and HapMap 3 SNPs excluding the MHC region as regression SNPs. We performed this analysis using summary statistics from 438 traits in UK Biobank (downloaded from https://nealelab.github.io/UKBB_ldsc) with highly statistically significant SNP heritability (LDSC z-score > 7, the threshold recommended in [91]).

### Expression variability across species

To understand the variability in expression between humans and other species, we focused on gene expression differences between human and chimpanzee as estimated from RNA sequencing of an *in vitro* model of the developing cerebral cortex for each species [42]. As a metric of variability between the two species, we used the absolute log-fold change (LFC) in gene expression between human and chimpanzee cortical spheroids, which was calculated from samples collected at several time points throughout differentiation of the spheroids. LFC estimates were obtained from Supplementary Table 9 of [42].

To visualize the relationship between constraint and absolute LFC, we plotted a LOESS curve between the constraint on a gene (gene rank from least to most constrained using either *s*_het_ or LOEUF as the constraint metric) and the absolute LFC for the gene. Curves were calculated using the `LOWESS` function from the `statsmodels` package with parameters `frac` = `0.15` and `delta` = `10`.

### Expression variability across individuals

To calculate a measure of expression variance across GTEx samples, we log-transformed the pergene mean and variance of gene expression levels (where expression is in units of Transcripts Per Million) and used the residuals from LOESS regression of the transformed expression variance on the transformed mean expression. LOESS regression was computed using the `LOWESS` function from the `statsmodels` package with parameters `frac` = `0.1` and `delta` = `0`. This procedure reduces the correlation between mean expression and expression variance (Spearman *ρ* = 0.02 between mean expression and residual variance, compared to Spearman *ρ* = 0.90 between mean expression and variance before regression). We calculated expression variance using 17,398 RNA-seq samples in the GTEx v8 release [43] (838 donors and 52 tissues/cell lines) for all genes with a median TPM of *≥* 5. LOESS curves for visualization were computed as in “Expression variability across species.”

## Feature interpretation

### Training models on feature subsets

We grouped features into categories (see Supplementary Table 4 for the features in each category), and trained a model for each category to predict *s*_het_ from the corresponding features. For each model, we tuned hyperparameters over a subset of the values we considered for the full model (Table 5), and chose the combination of hyperparameters that minimized the loss over genes in the validation set. As a baseline, we trained a model with no features, such that all genes have a shared prior distribution that is learned from the LOF data—this model is analogous to a standard empirical Bayes model.

**Table 5:** Parameters for feature subsets.

| Parameter(s) | Tested values |
| --- | --- |
| Learning rate | 0.01, 0.04 |
| Maximum tree depth (max_depth) | 3 |
| Data subsampling ratio (subsample) | 0.8, 1 |
| Minimum weight of a leaf node (min_child_weight) | 2, 4 |
| L1 regularization (alpha) | 1, 2 |
| L2 regularization (lambda) | 0 |
| Number of trees to fit per iteration (n_estimators) | 1 |

### Definition of expression feature subsets

We grouped gene expression features into 24 categories representing tissues, cell types, and developmental stage using terms present in the feature names (Table 6).

**Table 6:** Terms used to define tissues for expression features.

| Category | Terms in the feature (not case sensitive) |
| --- | --- |
| Brain | brain, nerve, microglia, hippocampus |
| Digestive | digestive, gut, gutendoderm, intestine, colon, ileum |
| Development | development, gastrulation, embryo |
| Lung | lung, airway |
| Eye | eye, retina |
| Endothelium | endothelium |
| Muscle | muscle |
| Hair follicle | hairfollicle |
| Kidney | kidney |
| Immune | immune, monocytes, nk, tcell, pbmc |
| Prostate | prostate |
| Blood | blood, heme, fetalblood |
| Adipocyte | adipocyte |
| Heart | heart, aorta |
| Thymus | thymus |
| Pancreas | pancreas, islets, pancreasductal |
| Liver | liver |
| Testis | testis |
| Synovial fibroblast | synovialfibroblast |
| Bladder | bladder |
| Placenta | placenta |
| Bone marrow | bonemarrow |
| CSF | csf |
| Lymph nodes | lymphnodes |

### Scoring individual features

To score individual gene features, we varied the value of one feature at a time and calculated the variance in predicted *s*_het_ as a feature score. In more detail, we fixed each feature to values spanning the range of observed values for that feature (0th, 2nd, …, 98th, and 100th percentile), such that all genes shared the same feature value. Then, for each of these 51 feature values, we averaged the *s*_het_ values predicted by the learned priors over all genes, where the predicted *s*_het_ for each gene is the mean of its prior. We denote this averaged prediction by 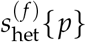 for some feature *f* and percentile *p*. Finally, we define the score for feature *f* as score 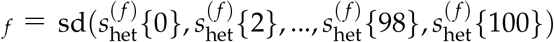, where sd is a function computing the sample standard deviation. In other words, a feature with a high score is one for which varying its value causes high variance in the predicted *s*_het_.

For the lineplots in Figures 4**C**-4**F**, we scale the predictions 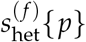 for each feature *f* by subtracting 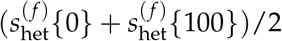 from each prediction.

### Pruning features before computing feature scores

While investigating the effects of features on predicted *s*_het_, we found that including highly correlated features in the model could produce unintuitive results, such as opposite correlations with *s*_het_ for highly similar features. Therefore, for Figures 4**C**-4**F**, we first pruned the set of features to minimize pairwise correlations between the remaining features. To do this, we randomly kept one feature in each group of correlated features, where such a group is defined as a set of features where each feature in the set has an absolute Spearman *ρ >* 0.7 to some other feature in the set.

For Figures 4**C**-4**F**, we trained models on the relevant features in this pruned set (gene ontology, network, gene regulatory, and gene structure features for Figures 4**C**, 4**D**, 4**E**, and 4**F** respectively).

## Supplementary Material

### A Empirical Bayes with NGBoost

#### Empirical Bayes overview

In the simplest version of empirical Bayes, we specify the form of the prior distribution and assume that that prior is shared across all genes—for example, for gene *i* we might assume the prior distribution is 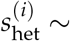 LogitNormal(*µ, σ*) with density 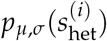, where the LogitNormal(*µ, σ*) distribution is defined such that logit 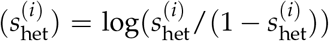 is normally distributed with mean *µ* and variance *σ*^2^. We can then estimate *µ* and *σ* using the observed LOF data for each gene, ***y***_1_, …, ***y***_*M*_, by maximizing the marginal likelihood:

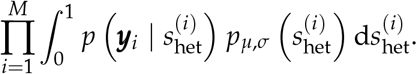

Next, we can compute the posterior distribution of 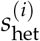 for each gene,

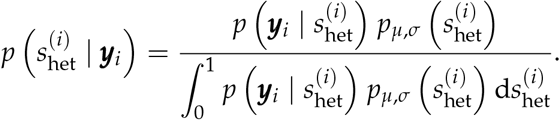

However, rather than learning the parameters for the prior from only the LOF data, we can also use gene features to learn gene-specific prior parameters, *µ*_*i*_ and *σ*_*i*_. To do this, we used a machine learning approach, `NGBoost`, to learn functions *f* and *g* such that *µ*_*i*_ = *f* (***x***_*i*_) and *σ*_*i*_ = *g*(***x***_*i*_), where ***x***_*i*_ is a vector of gene features associated with gene *i*. In the next few sections, we will describe how we learned *f* and *g*.

#### NGBoost

`NGBoost` (Natural Gradient Boosting) is an approach for training gradient boosted trees to predict the parameters of a probability distribution [16]. Gradient boosted trees are a type of machine learning model typically used to predict outcomes *y*, from features *X*, producing point estimates such as predictions of 𝔼[*y* | *X*]; in contrast, `NGBoost` uses gradient boosted trees to predict *p*(*y* | *X* = ***x***) by learning parameters of *p*(*y* | *X* = ***x***) as functions of ***x***—in other words, `NGBoost` allows us to learn the full distribution of *y* conditioned on observing the features ***x***.

Specifically, for gene *i*, we assume the prior distribution is 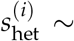 LogitNormal(*µ*_*i*_, *σ*_*i*_), with density 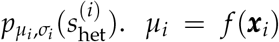 and *σ*_*i*_ = *g*(***x***_*i*_) are functions of the vector of gene features ***x***_*i*_, where *f* and *g* are parameterized as gradient-boosted trees. We chose this distribution as previous work has suggested that 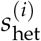 is distributed on a logarithmic scale [1, 2, 4], yet, 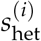 is also bounded between 0 and 1. Both of these properties are enforced by the LogitNormal distribution. In Supplementary Note B, we develop a population genetic likelihood 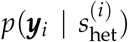, where ***y***_*i*_ is a vector that represents the observed frequencies of each possible loss of function variant for the gene. Then, with *M* genes in the training set, the score that `NGBoost` minimizes during training is:

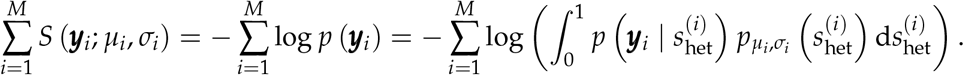

To do this, `NGBoost` first initializes the parameters of *f* and *g* such that all genes have the same prior distribution. Next, `NGBoost` adopts a gradient descent approach to minimize the score function: for each iteration until training ends, `NGBoost` first computes the gradient of gene *i*’s score with respect to the parameters *µ*_*i*_ and *σ*_*i*_ of 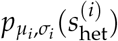. In the original implementation, `NGBoost` uses *natural gradients*, which take into account the underlying “information geometry” of the space of distributions in a way that standard gradients do not [92], but natural gradients are costly to compute, so we use standard gradients instead. After computing the gradient, `NGBoost` fits a decision tree to each dimension of the gradient, updating *µ*_*i*_ and *σ*_*i*_ in the direction that most steeply decreases the gene’s score. While gradient-boosting algorithms (including `NGBoost`, by default) typically fit a single decision tree at each iteration, we allow `NGBoost` to fit up to `n_estimators` trees per iteration, where `n_estimators` is a hyperparameter that we tune.

Below, we summarize the training algorithm. Let 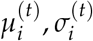 denote the parameters of the prior at training iteration *t*.

1. Initialize parameters for all genes, *i* = 1, …, *M*:

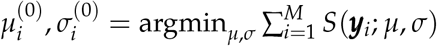
2. For iterations *t* = 1, …, *T*:
  a. For each gene, calculate gradients of the score: 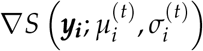, whose two components we denote as 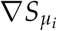 and 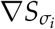
  b. Fit decision trees *f* ^(*t*)^ and *g*^(*t*)^ on the gradients:

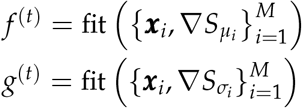
  c. Perform a line search to find a scaling, *ϕ*^(*t*)^ that optimizes the loss function along the search direction implied by *f* ^(*t*)^ and *g*^(*t*)^. That is, set:

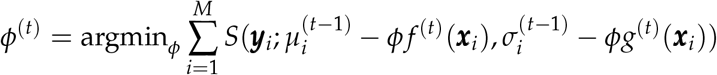

In practice, `NGBoost` approximately solves this optimization problem by initializing at *ϕ* = 1, iteratively doubling *ϕ* until the objective begins to increase, and then restarting at *ϕ* = 1 and iteratively halving *ϕ* until the objective begins to increase. Whichever of these *ϕ* minimized the objective function is used for *ϕ*^(*t*)^.
  d. Update the parameters for each gene, where *η* is a learning rate that is chosen by the user as a hyperparameter:

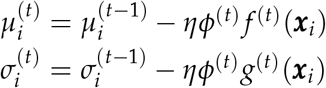

Once training is complete, we obtain a learned prior with parameters 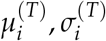, and can compute the posterior distribution of *s*_het_

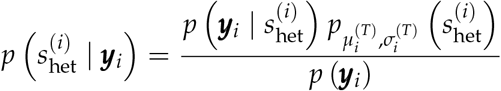

as well as the mean of this distribution

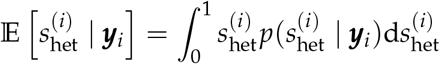

To compute 95% Credible Intervals, we compute the CDF of the posterior distribution using Pytorch’s `cumulative_trapezoid` function [93]. Then, the 95% Credible Interval per gene is defined as [lb^(*i*)^, ub^(*i*)^] such that 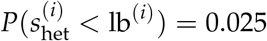 and 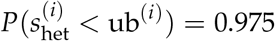.

#### NGBoost— implementation details

To initialize parameters (step 1 in the training algorithm), we perform gradient descent with the AdamW optimizer [94] implemented in `PyTorch` [93] with a learning rate of 1 *×* 10^*−*3^ and otherwise default settings. We initialize the optimization at *µ* = 0 and *σ* = 1.

We numerically compute all integrals using the trapezoid method implemented in `PyTorch`, which enables gradient computation through automatic differentiation. We perform all integrals using the transformed parameter, Logit(*s*_het_). Since *s*_het_ has a LogitNormal prior, this transformation has a Normal distribution, and hence puts all but a negligible fraction of its mass within 8 standard deviations, *σ*, of its mean, *µ*. Therefore, we perform the trapezoid method on an evenly spaced grid of 10^4^ points on the domain *µ −* 8*σ, µ* + 8*σ*.

To flexibly fit decision trees at each training iteration, we use the `XGBoost` package, a library used for fitting standard gradient boosted trees [95]. In comparison to the default `NGBoost` learner, `XGBoost` supports missing features and allows for adjustment of numerous hyperparameters (see “Training and Validation” in Methods). In contrast to typical applications of `XGBoost`, we only allow a few (`n_estimators` in Table 4) trees to be fit at each training iteration, as we are using `XGBoost` within a training loop rather than as a standalone approach for model fitting.

All distributions were implemented using `PyTorch`, and training was conducted with GPU support when available, with `tree_method = “gpu_hist”` for the `XGBoost` learners.

### B Population Genetics Model

#### Overview of model

Some of the most commonly used measures of gene constraint (pLI [11], LOEUF [12]) are framed in terms of the number of unique LOFs observed in gene, *O*, relative to the number expected under a null model, *E*. While operationalizing constraint as some function of *O* and *E* captures the intuition that seeing fewer LOFs than expected is evidence that a gene is conserved, the numerical values of pLI and LOEUF are difficult to interpret. In practice this means that such measures can be useful for ranking which genes are important, but it makes it difficult to contextualize these results in terms of other types of variants, such as missense or noncoding variants, or copy number variants. Previous approaches have pioneered using a population genetics model in this context to obtain interpretable estimates, albeit with different technical details that we discuss below [1, 2, 4].

In order to obtain a more interpretable measure of constraint, we formalize constraint as the strength of natural selection acting against gene loss-of-function in a population genetics model. That is, we can ask how much fitness is reduced on average for an individual with one or two non-functional copies of a gene relative to individuals with two functional copies, following previous work [1, 2, 4]. To tie this concept of constraint to observed allele frequency data, we use a slightly simplified version of the discrete-time Wright Fisher model. This model contains mutation, selection, and genetic drift, and assumes that there are only two alleles and that the population is panmictic, monoecious, and has non-overlapping generations. While all of these assumptions are violated in humans (there are four nucleotides, population structure, two sexes, and overlapping generations), the model still provides a good approximation to allele frequency dynamics through time. If the allele frequency in generation *k* is *f*_*k*_, then we model the allele frequency in the next generation via binomial sampling:

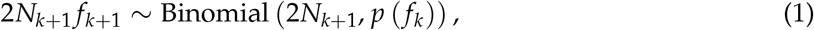

where *N*_*k*+1_ is the number of diploid individuals in generation *k* + 1, with

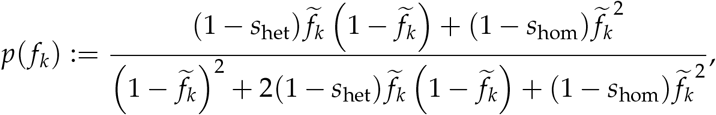

where 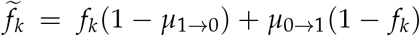 is the allele frequency after alleles change from non-LOF to LOF at rate *µ*_0*→*1_ and from LOF to non-LOF at rate *µ*_1*→*0_. The function *p*(*·*) arises from considering bidirectional mutation and approximating a model of diploid selection where the relative reproductive success of individuals with 0, 1, or 2 copies of the LOF are 1, 1 *− s*_het_, and 1 *− s*_hom_ respectively [13]. In practice, most LOF variants are extremely rare, and so it is exceedingly unlikely to find individuals homozygous for the LOF. This makes estimating *s*_hom_ as a separate parameter very difficult, and so we instead assume that *s*_hom_ = min *{*2*s*_het_, 1*}*. This is equivalent to assuming genic selection (i.e., additive fitness effects) with the constraint that an individual’s relative fitness cannot be lower than 0.

Equation 1 fully specifies the model except for an initial condition. That is, we need to know what the distribution of frequencies is in generation 0. One mathematically appealing choice would be to assume that the population is at equilibrium at time 0, but this seemingly straight-forward choice results in nonsensical conclusions. To see why, if the mutation rates are low and selection is negligible, then at equilibrium, with extremely high probability the population will either be in a state where the frequency of the LOF allele is very close to zero or in a state where the frequency of the LOF allele is very close to one. If the mutation rates between the two alleles are close to equal, then these two cases happen roughly equally often. That is, we would expect there to be a ~50% chance that the population is fixed or nearly fixed for the LOF mutation. If there are multiple independently evolving sites at which a LOF could arise (or if there are many more ways to mutate to a LOF state than a non-LOF state), then the chance that any of these sites is fixed or nearly fixed for a LOF rapidly approaches 10 0%. Under this equilibrium assumption, we thus reach the absurd conclusion that the mere act of observing a gene that is functional in a majority of the population is overwhelming evidence that the gene is strongly selected for. Another way of viewing this is that in reality we can only observe genes that are functional in an appreciable fraction of the population, and so we should somehow be conditioning on this event, whereas the equilibrium assumption looks at a given randomly chosen stretch of DNA and asks whether it could be a gene given some set of mutations. Indeed, any randomly chosen stretch of DNA could be made a gene through a series of mutations, but for any given stretch it would be extremely unlikely to be a functional gene, and the equilibrium assumption exactly captures how rare this would be.

We instead use the equilibrium of another process as the initial condition, which avoids these conceptual pitfalls. We assume the distribution of frequencies at generation 0 is the equilibrium conditioned on the LOF allele never reaching fixation in the population. We then compute the likelihood of observing a given present-day frequency while continuing to condition on non-fixation of the LOF allele. This assumption implies that no matter the current frequency of the LOF variant, we know that at some point in the past the population was fixed for the functional version of the gene, and the LOF variant can thus be thought of as being “derived” and the non-LOF variant “ancestral”. In the limit of infinitely low (but non-zero) mutation rates, this assumption becomes equivalent to the commonly assumed “infinite sites” model commonly used to compute frequency in population genetics [96]. In contrast to the infinite sites model, where the probability that any given site is segregating must be 0, our model allows us to compute the probability that a given site is segregating. Furthermore, we can easily model recurrent mutation which can be important for sites with large mutation rates (such as CpGs) and large sample sizes [97], whereas under the infinite sites model each mutation necessarily happens at a unique position in the genome, ruling out the possibility of recurrent mutation. Below, we will write *p*_DTWF_(*y* | *s*_het_) for the probability mass function computed using this procedure, with “DTWF” representing Discrete-Time Wright-Fisher, and *y* being an observed LOF allele frequency.

Equation 1 is easy to describe and simulate under, and a very similar model has been used in an approximate Bayesian computation approach to estimate *s*_het_ [4]. While simulation is easy, computing likelihoods under this model is difficult for large sample sizes, and unfortunately we need explicit likelihoods in our empirical Bayes approach. In recent work [15], we have developed an efficient method for computing likelihoods under this model. The key idea is that the above dynamics can be written as

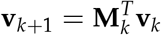

where **v**_*k*_ is a vector of dimension 2*N* + 1 where entry *i* is the probability that there are *i* haploids that have the LOF allele in generation *k*, and **M**_*k*_ is a matrix where row *i* is the the probability mass function of the Binomial distribution in Equation 1 given that the allele frequency in generation *k* is *i*/2*N*_*k*_. This formulation makes clear that we can obtain the likelihood of observing a given frequency at present given some initial distribution by performing a series of matrix-vector multiplications. Naively this would be prohibitively slow as **M**_*k*_ can be as large as 10^7^ *×* 10^7^, but in [15] we show that **M**_*k*_ is approximately highly structured — it is both approximately extremely sparse and approximately extremely low rank. Combining these insights we can perform matrix-vector multiplication that is provably accurate while reducing the runtime for matrix-vector multiplication from 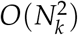 to *O*(*N*_*k*_). Similar insights can be used to speed up the computation of equilibria, which we discuss in detail in [15]. Furthermore, as discussed above, we actually want to compute likelihoods conditioned on non-fixation of the LOF allele, but that is as simple as setting the column of **M**_*k*_ corresponding to fixation to 0, and then renormalizing **v**. We precompute these likelihoods for each possible pair of mutation rates (to and from the LOF allele) across a range of *s*_het_ values (100 log-linearly spaced points between 10^*−*8^ and 1, as well as 0). We describe how we set the mutation rates and the population sizes implicit in **M**_*k*_ below.

#### Modeling misannotation of LOFs

Under the likelihood described above, and as seen in Figure 2**A**, positions where a LOF variant could occur, but no LOF alleles are observed are slight evidence in favor of selection, while high frequency variants are extremely strong evidence against selection. Meanwhile, we suspect that many variants that are annotated as causing LOF actually have little to no effect on the gene product due to some form of misannotation. If these misannotated variants evolve effectively neutrally, they can reach high frequencies and cause us to artifactually infer artificially low levels of selection. These misannotated variants can be particularly problematic for approaches that combine frequencies across all LOFs within a gene to obtain an aggregate gene-level LOF frequency [1,2,4].

LOEUF [12] and pLI [11] avoid this problem by throwing away all frequency information except for whether a LOF is segregating or not. While this approach is more robust, the ignored frequency information is extremely useful for estimating the strength of selection. For example, consider a gene where we expect to see 5 unique LOFs under neutrality and we see 3 segregating LOFs. This might seem like weak or negligible constraint (*O*/*E* = 0.6), but if those 3 sites are all highly mutable and the variants at those sites are each only present in a single individual, then it is plausible that this gene is quite constrained.

To take full advantage of the information in the LOF frequencies while remaining robust to misannotation, we take a composite likelihood approach [98], closely related to the Poisson random field assumption commonly used in population genetics [96]. We approximate gene-level likelihoods as a product of variant level likelihoods

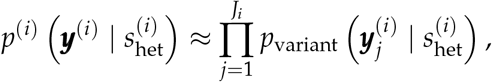

where ***y***^(*i*)^ is a vector of the observed allele frequencies at each possible LOF site in gene *i*, and 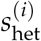 is the selection coefficient for having a heterozygous loss-of-function of gene *i*. Under this formulation, we can easily model misannotation by assuming that each LOF independently has some probability of being misannotated, *p*_miss_, and that misannotated variants evolve neutrally:

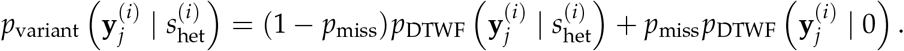

Using this formulation, we can take full advantage of the rich information included in the exact sample frequencies of each LOF variant, while still being robust to occasional misannotation. In practice, we precompute *p*_variant_ using a grid of *p*_miss_ values, and then to obtain the likelihood at arbitrary values of *s*_het_ and *p*_miss_ we linearly interpolate in log-likelihood space. Below, we discuss our approach for setting *p*_miss_.

Given a probability of misannoation, we can then calculate a posterior probability that any given variant has been misannotated. We include a table of these misannotation probabilities for all possible LOFs in [80].

Supplementary Figure 2 shows the impact of modeling misannotation on the variant-level likelihoods. In particular we present the likelihood curves for individual variants of different frequencies at a high mutation rate site (analogous to Figure 2**A**). We see that increasing *p*_miss_ affects the likelihoods in two ways. First, it slightly flattens the overall likelihood, which makes sense as some information must be lost as we assume more and more LOFs are misannotated. Second, including *p*_miss_ results in a floor on how low the likelihood can go, and higher values of *p*_miss_ result in higher floors. This also makes sense, as any individual variant can always be “explained away” by it being misannotated, and in particular, the likelihood can never be lower than *p*_miss_ times the likelihood of the variant under neutrality.

Supplementary Figure 3 shows the impact of *p*_miss_ on gene-level likelihoods for two example genes. For *AARD*, a gene with only 4.3 expected LOFs and only one observed LOF, *p*_miss_ has a large impact on the log-likelihood. For low values of *p*_miss_ the single segregating LOF (present at a frequency of 42/250,000) provides enough evidence to strongly rule out *s*_het_ *>* 0.02, with the likelihood rapidly approaching zero. In contrast, if there were no LOFs, then the likelihood would monotonically increase to a maximum at *s*_het_ = 1. As such, as *p*_miss_ increases, we start to see bimodal likelihoods, due to being a mixture of the case where the segregating variant is not misannotated (where the likelihood would be monotonically decreasing), and the case where the segregating variant is misannotated (where the likelihood would be monotonically increasing). For *LPA*, which has 90.2 expected LOFs and 118 observed LOFs, there are enough variants of similar frequencies where even for *p*_miss_ as high as 0.1 it is unlikely for all of the LOFs to be due to misannotation. As a result, the likelihood is relatively insensitive to *p*_miss_, with *p*_miss_ primarily acting to slightly flatten the likelihood.

We also investigated whether the inferred posterior probabilities of misannotation vary based on their positions within transcripts. We mapped each LOF to its position relative to the start and end of the canonical transcript for each gene. We found that LOFs due to early stop codons are inferred to be more likely to be misannotated if they are near the start or end of transcripts, possibly due to alternative translation initiation [99] or avoiding the nonsense-mediated decay pathway [100] respectively. In contrast, variants that are annotated as LOFs due to having a predicted effect on splicing are increasingly likely to be misannotated toward the end of transcripts. These results are presented in Supplementary Figure 4.

As an example of the importance of correcting for misannotation, we consider the case of the gene *PPFIA3* (ENSG00000177380). This gene has a LOEUF score of 0.12 and so appears very constrained, but in an early version of our model where we did not incorporate variant misannotation, we inferred a posterior mean value of *s*_het_ of ~2 *×* 10^*−*4^, which is right at the border of being nearly neutral. Inspecting the LOF data for this gene, we find that all potential LOFs are either not observed or observed in a single individual, except for a single splice donor-disrupting variant at 16% frequency. There are no obvious signs indicating that this variant is misannotated (e.g., in terms of coverage or mappability). If we model misannotation, however, we find that this variant is likely misannotated (posterior probability of misannotation ~100%), and as a result we estimate extremely strong selection against gene loss-of-function (posterior mean *s*_het_ of ~ 0.202). Indeed, a single autosomal dominant missense variant in this gene is suspected to have caused a number of severe symptoms including developmental delay, intellectual disability, seizures, and macrocephaly in an Undiagnosed Diseases Network participant (https://undiagnosed.hms.harvard.edu/participants/participant-159/) [101].

#### Modeling the X chromosome

We must slightly modify our model when applying it to the X chromosome. Because males only have one copy of the X chromosome, there are only 3/4 as many X chromosomes as autosomes (assuming an approximately equal sex ratio). As a result, when dealing with the X chromosome we scale all population sizes to 3/4 of the size used for the autosomes (rounded to the nearest integer). We also need to slightly modify the expected frequency in the next generation. We assume haploid selection in males with strength *s*_hom_, and diploid selection in females with selection coefficients *s*_het_ and *s*_hom_ for individuals heterozygous and homozygous for the LOF variant respectively. This selection results in modified allele frequencies in the pool of males and females, and the we assume that each chromosome in the next generation has 1/3 probability of coming from a male, and 2/3 probability of coming from a female. This means that the expected frequency in the next generation is 1/3 times the post-selection frequency in males plus 2/3 times the post-selection frequency in females. Variants within the pseudoautosomal regions on the X are modeled identically to variants on the autosomes. Agarwal and colleagues also considered selection on the X in the context of LOF variants, with a model similar to that described here [4].

#### Model parameters

Our model has three key parameters — the mutation rate, the demographic model (i.e., population sizes through time), and the probability that different variants are misannotated.

We obtained mutation rates from gnomAD [12, Supplementary Dataset 10], which take into account trinucleotide context and methylation level (for CpG to TpG mutations). In our population genetics model, we assume that there are only two alleles (a functional allele and a LOF allele), whereas in reality there are four nucleotides. We approximate the rate of mutating from the functional allele to the LOF allele as being the sum of the mutation rates from the reference nucleotide to any nucleotide that might result in LOF. For example, if the reference allele is A, and either a C or a T would result in LOF, then we say that the rate at which the functional allele mutates to the LOF allele is the rate at which A mutates to C in this context plus the rate at which A mutates to T in this context. For the rate of back mutation from the LOF allele to the functional allele, we compute a weighted average of the rates of each possible LOF nucleotide back-mutating to any possible non-LOF nucleotide, weighed by the probability that the original non-LOF nucleotide mutated to that particular LOF nucleotide. Continuing our previous example, suppose A mutates to C at rate 1 *×* 10^*−*8^ and A mutates to T at a rate 1.5 *×* 10^*−*8^. Then conditioned on there having been a single mutation resulting in a LOF variant, there is a 1/2.5 = 0.4 chance that the LOF is C and 0.6 chance that the LOF is T. We then compute the back mutation rate as 0.4 times the rate at which C mutates to A in this context plus the rate at which C mutates to G in this context (since both A and G do not result in LOF) plus 0.6 times the rate at which T mutates to A in this context plus the rate at which T mutates to G in this context. Implicitly this scheme assumes that the flanking nucleotides in the trinucleotide context do not change, and we further assume that all mutations resulting in CpGs result in unmethylated CpGs.

For the population sizes in each generation, we used the “CEU” model inferred in [81] using the 1000 Genomes Project data [102]. This model was also used in [4]. Population sizes under this model are relatively constant before 5156 generations ago (approximately 155 thousand years ago) and the effects of strong selection are relatively insensitive to all but the most recent population sizes, so for a computational speedup we assumed that the population size was constant prior to 5156 generations ago. Recently, [4] found that this CEU model underestimates the number of low frequency variants and that changing the population size to 5,000,000 for the most recent 50 generations provides a better fit to the data. We used both demographic models and found qualitatively similar results, with slightly better fit provided by the modified model, so we used that demographic model for all subsequent analyses. In both cases, we modified the most ancient population sizes, which are relatively constant, to be actually constant to speed up likelihood calculations. The demographic models are presented in Supplementary Figure 5.

Given that demography plays an important role in the likelihood and that gnomAD contains individuals of diverse ancestries, we wanted to make sure that our results were generally robust to misspecification of the demography. All of the results presented in the main text used the entirety of gnomAD v2, but we also trained models using the subset of individuals labeled in the dataset as “non-Finnish European” (NFE) as well as all non-NFE individuals. When training these models we assumed the Agarwal. *et al*. demography, regardless of the ancestry of the individuals used in training. The posterior mean *s*_het_ values under all three models are quantitatively and qualitatively consistent, with Spearman correlations greater than 0.93 between all pairs Figure 6. The high concordance indicates that our pooling of individuals of different genetic ancestries is justified, and that our results are robust to slight mismatches between the demography and the individuals used.

The only remaining model parameter is *p*_miss_ the probability that any given LOF is misannotated. Throughout we focus on LOFs that either introduce early stop codons, disrupt splice donors, or disrupts splice acceptors. Given that predicting which variants have these different consequences involves different bioinformatic challenges, we inferred separate misannoatation probabilities 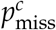 for *c ∈ {*stop codon, splice donor, splice acceptor*}*. Below we write *p*_miss_ for the collection of these three misannotation parameters. To get a rough estimate of these parameters and avoid excessive computational burden, we took an h-likelihood approach [103, 104]. That is, we jointly maximized the likelihood across all genes with respect to their selective constraints as well as the the three misannotation probabilities that are shared across all genes:

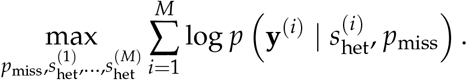

This approach of just using the maximum likelihood estimates of *s*_het_ for each gene contrasts with the standard empirical Bayes approach, which would involve marginalizing out the unknown *s*_het_ values. Yet, this marginalization step depends on the prior on *s*_het_, which we learn via our `NGBoost` framework. As a result, we would need to repeatedly run our `NGBoost` framework as an inner loop to perform the standard empirical Bayes approach on *p*_miss_. For our application, these values are nuisance parameters, and the results are relatively insensitive to their exact values so we opted for this simpler h-likelihood approach. Ultimately, we estimate that the probability of misannotation is 0.7%, 4.5%, and 8.1% for stop codons, splice donors, and splice acceptors respectively.

### C Feature processing and selection

We compiled 10 types of gene features from several sources:

1. *Gene structure*. Gene structure features were derived from GENCODE gene annotations (Release 39) [84]. Such features include the number of transcripts and, for the primary transcript of each gene (the transcript tagged `Ensembl_canonical`), the number of exons as well as the length and GC content of the transcript, total coding region, 5’ UTR, and 3’ UTR.
2. *Gene expression*. We used gene features from 77 bulk and single-cell RNA-seq datasets, processed and derived in [105]. These datasets can be grouped into 24 categories representing tissues, cell types, and developmental stage (Table 6). For each dataset, features were derived separately from all data and from individual cell clusters (for example, gene loadings on principal components). In addition, features were derived from comparisons between clusters (for example, t-statistics for differential expression). Finally, we include a metric, *τ*, that summarizes the tissue-specificity of gene expression [106].
3. *Biological pathways and Gene Ontology terms*. First, we included previously curated biological pathway features [105, 107]. In addition, to include GO terms that capture additional known relationships between genes, we downloaded Biological Pathway (BP), Molecular Function (MF), and Cellular Component (CC) terms [108] with at least 10 member genes using the procedure described in [10]. Features for each gene were encoded as binary indicators of the gene’s membership in the pathways and GO terms.
4. *Connectedness in protein-protein interaction (PPI) networks*. We included previously computed measures of the connectedness of protein products of genes in PPI networks [10]. Connectedness was calculated as the number of interactions per protein weighted by the interaction confidence scores.
5. *Co-expression*. First, we included previously computed measures of the connectedness of genes in co-expression networks [10], where connectedness measures the relative number of neighbors of each gene in the network, averaged over tissues. Next, for each gene, we derived features representing its co-expression with other genes (i.e. correlation in their expression levels across samples). To do this, we downloaded from the GeneFriends database a co-expression network derived from GTEx RNA-seq samples [109,110], calculated the variance in the co-expression for each gene, and kept the 6,000 most variable genes. Then, we included the co-expression with each of these 6,000 genes as a feature.
6. *Gene regulatory landscape*. Gene regulatory features include the counts and properties of the enhancers and promoters that regulate each gene. First, we included the number of promoters per gene estimated by the FANTOM consortium using Cap Analysis of Gene Expression [10, 111]. Next, for each gene, we calculated the number, summed length, and summed score of enhancer-to-gene links predicted using the Activity-By-Contact (ABC) approach [49, 112], where an enhancer is considered linked to a gene if its ABC score is *≥*0.015. We computed separate features for each of 131 biosamples. We also included features derived by aggregating over all biosamples for both ABC enhancers and predicted enhancers from the Roadmap Epigenomics Consortium [10, 113, 114]—these feature include the number of biosamples with an active enhancer element, the total number of enhancer elements, the total number of enhancer elements after taking merging enhancer domains, the total length of the merged domains, and the average total enhancer length in an active cell type. Finally, we included the enhancer-domain score for each gene [9] as a feature.
7. *Conservation across species*. For each gene, we calculated the mean and 95th percentile phast-Cons scores over the gene’s exons for multiple alignments of 7, 17, 20, 30, and 100 vertebrate species to the human genome [115]. We downloaded phastCons Scores from https://hgdownload.soe.ucsc.edu/goldenPath/hg38/. In addition, we included the fraction of coding sequence (CDS) or exons constrained across 240 mammals or 43 primates sequenced in the Zoonomia project [116], with constraint determined by the per-base phyloP [117] or phastCons score. Zoonomia data were downloaded from https://figshare.com/articles/dataset/geneMatrix/13335548.
8. *Protein embedding features*. We included as features the embeddings learned by an autoencoder (ProtT5) trained on protein sequences [118]. Embeddings were downloaded from https://zenodo.org/record/5047020. The embedding for each protein is a fixed-size vector that captures some of the protein’s biophysical and functional properties. For each gene with more than one protein product, we averaged the embeddings of the proteins for that gene.
9. *Subcellular localization*. We included as features the subcellular localization of each protein and whether the protein is membrane-bound or soluble, as predicted by deep neural networks trained on the ProtT5 protein embeddings [118, 119]. Possible subcellular classes included nucleus, cytoplasm, extracellular space, mitochondrion, cell membrane, endoplasmatic reticulum, plastid, Golgi apparatus, lysosome or vacuole, and peroxisome. Predictions were one-hot encoded, and for each gene with more than one protein product, we summed the predictions for the gene’s proteins. Predictions were downloaded from https://zenodo.org/record/5047020.
10. *Missense constraint*. We included a measure of each gene’s average intolerance to missense variants (UNEECON-G score) [19]. UNEECON-G scores incorporate variant-level features to account for differences in the effects of missense variants on gene function.

In addition to these 10 groups of features, we included a binary indicator for whether the gene is located on the X chromosome. Genes in the pseudoautosomal regions were categorized as autosomal.

After compiling these features (total of 65,383), we performed feature selection to minimize the practical complexity of training on such a large feature set and the complexity of the resulting model. First, we removed features with zero variance and features where the Spearman correlation of the feature values with *O*/*E* (the ratio of observed over expected unique LOF variants, computed using gnomAD data) was less than 0.1 or had a nominal p-value *≥* 0.05. Next, we performed simultaneous feature selection and an initial round of hyperparameter tuning using the `shap-hypetune` package, which uses Bayesian optimization to identify a set of features and hyper-parameters that minimize the loss of a machine learning model fit on the training data. Specifically, we fit gradient-boosted trees using `XGBoost` to predict *O*/*E* from the gene features; we chose to perform feature selection using `XGBoost` rather than `NGBoost` as training `XGBoost` models is sub-stantially faster, and because we expect features/hyperparameters that perform well for `XGBoost` to also perform well for `NGBoost`. For each set of hyperparameters, `shap-hypetune` performs backward step-wise selection by removing the *k* least influential features (we chose *k* = 1000 and calculated influence using SHAP scores) at each step. Finally, we performed further feature selection using `shap-hypetune` by fixing the hyperparameters and performing backward step-wise selection with *k* = 50. Ultimately, we included 1,248 features in the model.

### D Potential for bias from the use of missense constraint or cross-species conservation

In theory, features that correlate with patterns of human LOF polymophism but not with constraint may bias our estimates. In particular, a number of population genetic forces beyond natural selection contribute to patterns of conservation across species and missense constraint, and if these forces also affect observed LOF frequencies, then these features could be problematic. While we take into account mutation rate differences due to trinucleotide context and methylation status when considering LOFs, larger-scale variation in mutation rates and other forces that might alter the “local effective population size” could affect various measures of constraint as well as LOF frequency in a manner independent of selection.

To evaluate this possibility, we trained a model excluding missense constraint and cross-species conservation features, and compared the *s*_het_ estimates from this model to those from the full model (Supplementary Figure 8). We find that the posterior mean values of *s*_het_ are highly correlated between the models (Supplementary Figure 8**A**, Spearman *ρ* = 0.92). In addition, the performance of the models in classifying genes essential *in vitro* and in classifying developmental disorder genes is extremely similar (Supplementary Figure 8**B,C**). These results indicate that the use of such features does not substantially bias the model.

### E Estimating additional gene properties using GeneBayes

GeneBayes is a flexible framework that can be used to infer other gene-level properties of interest beyond *s*_het_. In Figure 6, we presented a schematic of the key components of GeneBayes that users should specify, which we describe in more detail now.

First, users should specify the gene features to use as predictors. We expect the gene features we use for *s*_het_ estimation to work well for other applications, but GeneBayes supports any choice of features. In particular, GeneBayes can handle categorical and continuous features without feature scaling, as well as features with missing values.

Next, users should specify the form of the prior distribution. GeneBayes supports the distributions defined by the `distributions` package of `PyTorch`. GeneBayes also supports custom distributions, as long as they implement the methods used by GeneBayes (i.e. `log_prob` and `sample`) and are differentiable within the `PyTorch` framework.

Finally, users need to specify a likelihood function that relates their gene property of interest to observed data. The likelihood can be specified in terms of a `PyTorch` distribution, or as a custom function.

After model training, GeneBayes outputs a per-gene posterior mean and 95% credible interval for the property of interest. For each parameter in the prior, GeneBayes also outputs a metric for each feature that represents the contribution of the feature to predictions of the parameter.

In the next section, we describe in more detail the two example applications that we outlined in Figure 6.

#### Example applications

##### Differential expression

In this example, users have estimates of log-fold changes in gene expression between conditions and their standard errors from a differential expression workflow, and would like to estimate log-fold changes with greater power (e.g. for lowly-expressed genes with noisy estimates).

###### Likelihood

We define 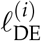 and *𝓁*_*i*_ as the estimated and true log-fold change in expression respectively for gene *i*, and *s*_*i*_ as the standard error for the estimate. Then, we define the likelihood for *𝓁*_*i*_ as

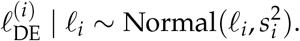

###### Prior

We describe two potential priors that one may choose to try. The first is a normal prior with parameters *µ*_*i*_ and *σ*_*i*_:

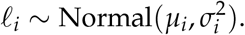

The second is a spike-and-slab prior with parameters *π*_*i*_, *µ*_*i*_, and *σ*_*i*_, which assumes that gene *I* only has a *π*_*i*_ probability of being differentially expressed:

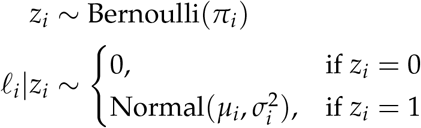

##### Variant burden tests

In this example, users have sequencing data from patients with a disease or (if calling *de novo* mutations) sequencing data from family trios, and would like to identify genes with excess mutational burden in patients (e.g. an excess of missense or LOF variants). One approach is to infer the relative risk for each gene (denoted as *γ*_*i*_ for gene *i*), defined as the expected ratio of the number of variants in patients to the number of variants in healthy individuals.

###### Likelihood

Let *E*_*i*_ be the number of variants we expect to observe for gene *i* given the study sample size and sequence-dependent mutation rates (e.g. expected counts obtained using the mutational model developed by [90]). Next, let *O*_*i*_ be the number of variants observed in patients for gene *i*. Then, we define the likelihood for *η*_*i*_ as

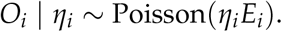

###### Prior

Because *η*_*i*_ is non-negative, one may want to choose a gamma prior with parameters *α*_*i*_ and *β*_*i*_:

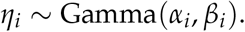

**Supplementary Table 1:** LOEUF and s_het_ for ribosomal proteins associated with Diamond-Blackfan anemia.

| Gene | $s_{\text{het}}$ | LOEUF | Obs. | Exp. |
| --- | --- | --- | --- | --- |
| *RPL18* | 0.74 | 0.28 | 0 | 10.5 |
| *RPL11* | 0.72 | 0.30 | 0 | 10.1 |
| *RPS19* | 0.70 | 0.37 | 0 | 8.1 |
| *RPS15A* | 0.68 | 0.56 | 0 | 5.4 |
| *RPL35A* | 0.56 | 0.41 | 0 | 7.3 |
| *RPS26* | 0.56 | 0.48 | 0 | 6.2 |
| *RPS7* | 0.56 | 0.31 | 0 | 9.7 |
| *RPL26* | 0.50 | 0.38 | 0 | 8.0 |
| *RPS24* | 0.49 | 0.59 | 1 | 8.0 |
| *RPL35* | 0.46 | 0.72 | 1 | 6.6 |
| *RPS10* | 0.45 | 0.27 | 0 | 11.0 |
| *RPL5* | 0.45 | 0.17 | 0 | 17.9 |
| *RPL27* | 0.43 | 0.48 | 0 | 6.2 |
| *RPL15* | 0.23 | 0.27 | 0 | 11.0 |
| *RPS29* | 0.19 | 1.2 | 1 | 3.9 |
| *RPS28* | 0.13 | 0.80 | 0 | 3.8 |
| *RPS27* | 0.10 | 0.64 | 0 | 4.7 |
| *RPS17* | 0.05 | 1.8 | 0 | 0.4 |

**Supplementary Figure 1:**
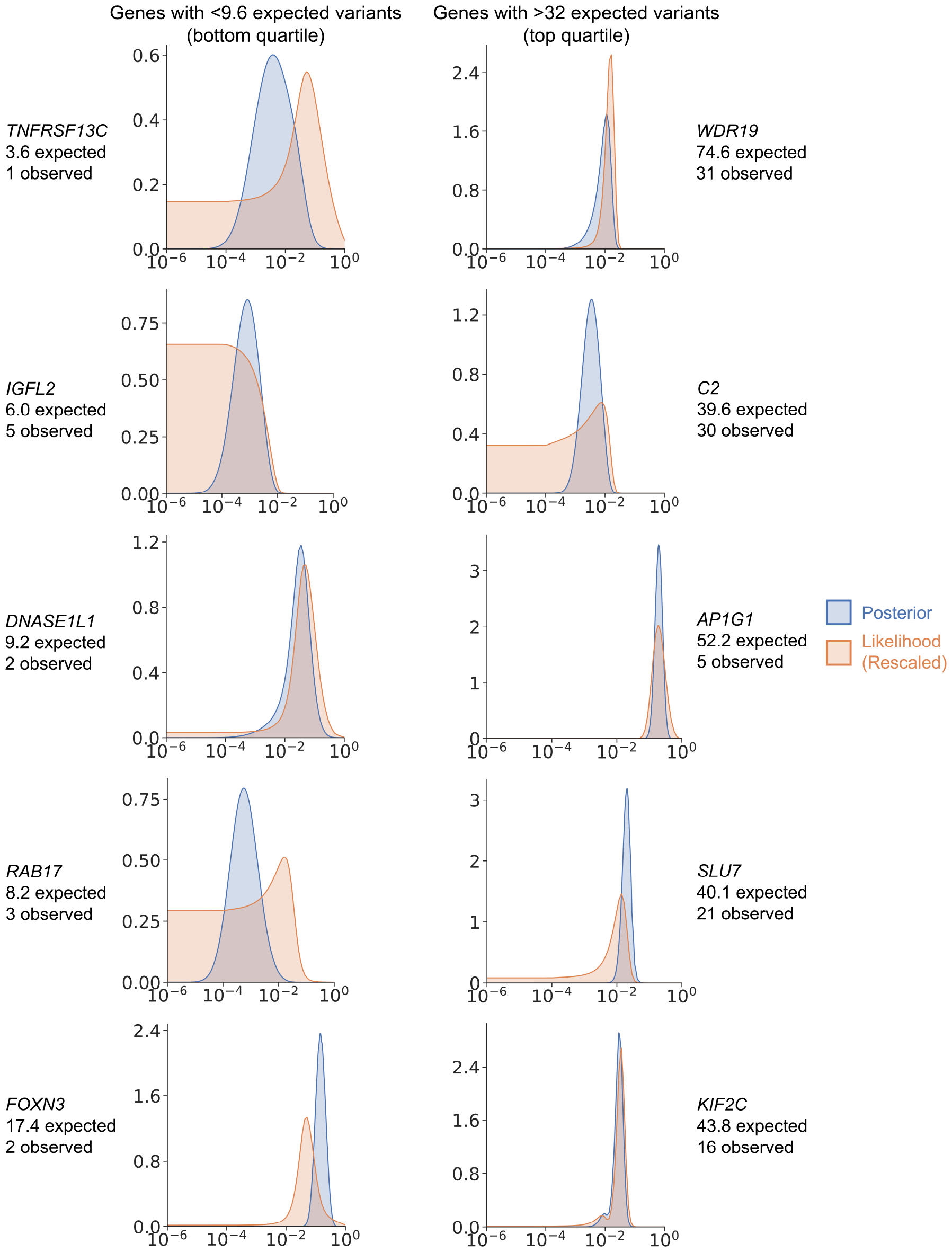
*s*_het_ distributions for additional example genes. Left: Posterior distributions and rescaled likelihoods for genes with few expected LOFs (genes in the bottom quartile). Right: Posterior distributions and rescaled likelihoods for genes with many expected LOFs (genes in the top quartile).

**Supplementary Figure 2:**
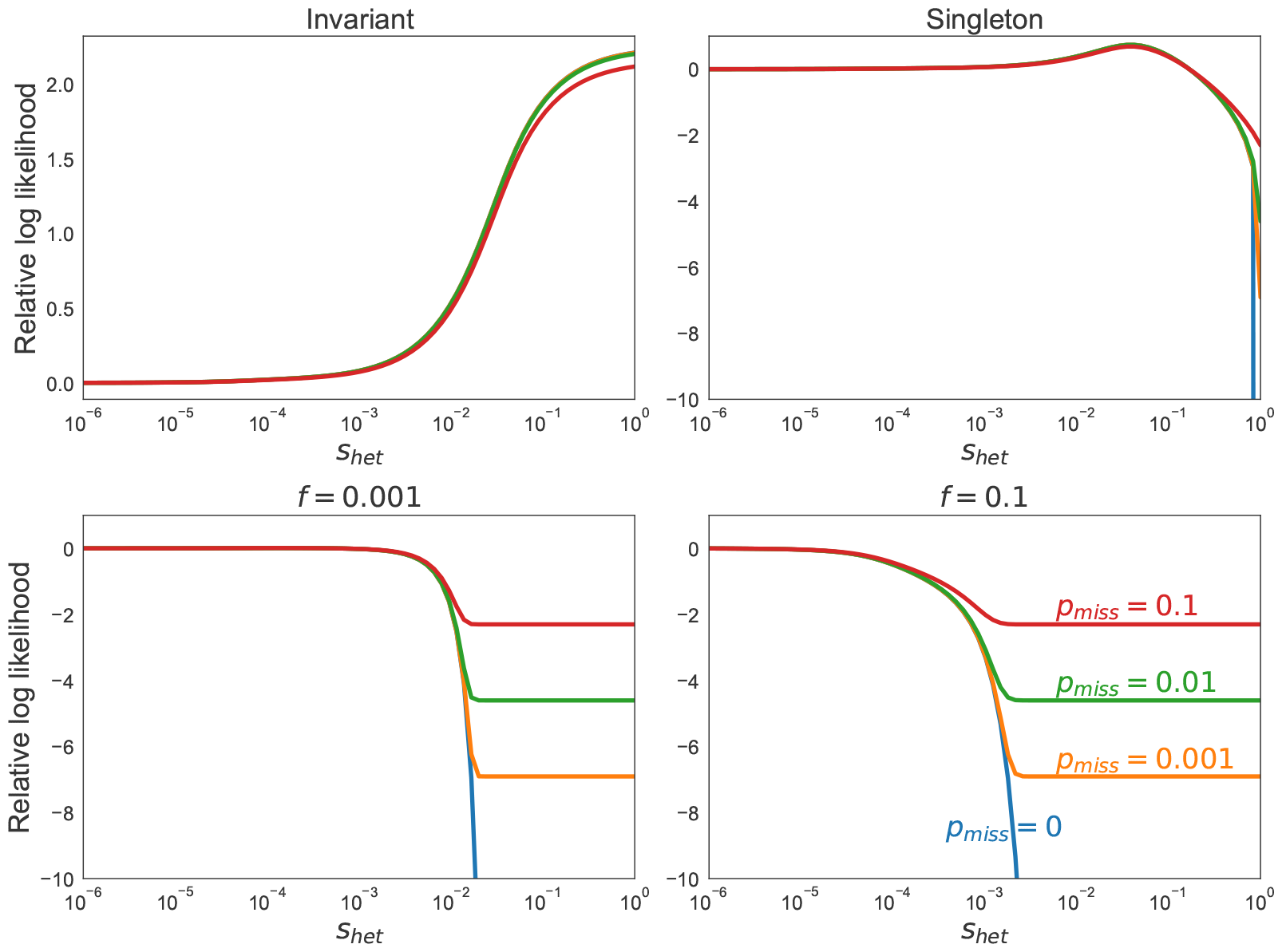
Dependence of likelihoods on parameters. Relative log likelihoods for a high mutation rate site with different allele frequencies, f, and misannotation probabilities, p_miss_.

**Supplementary Figure 3:**
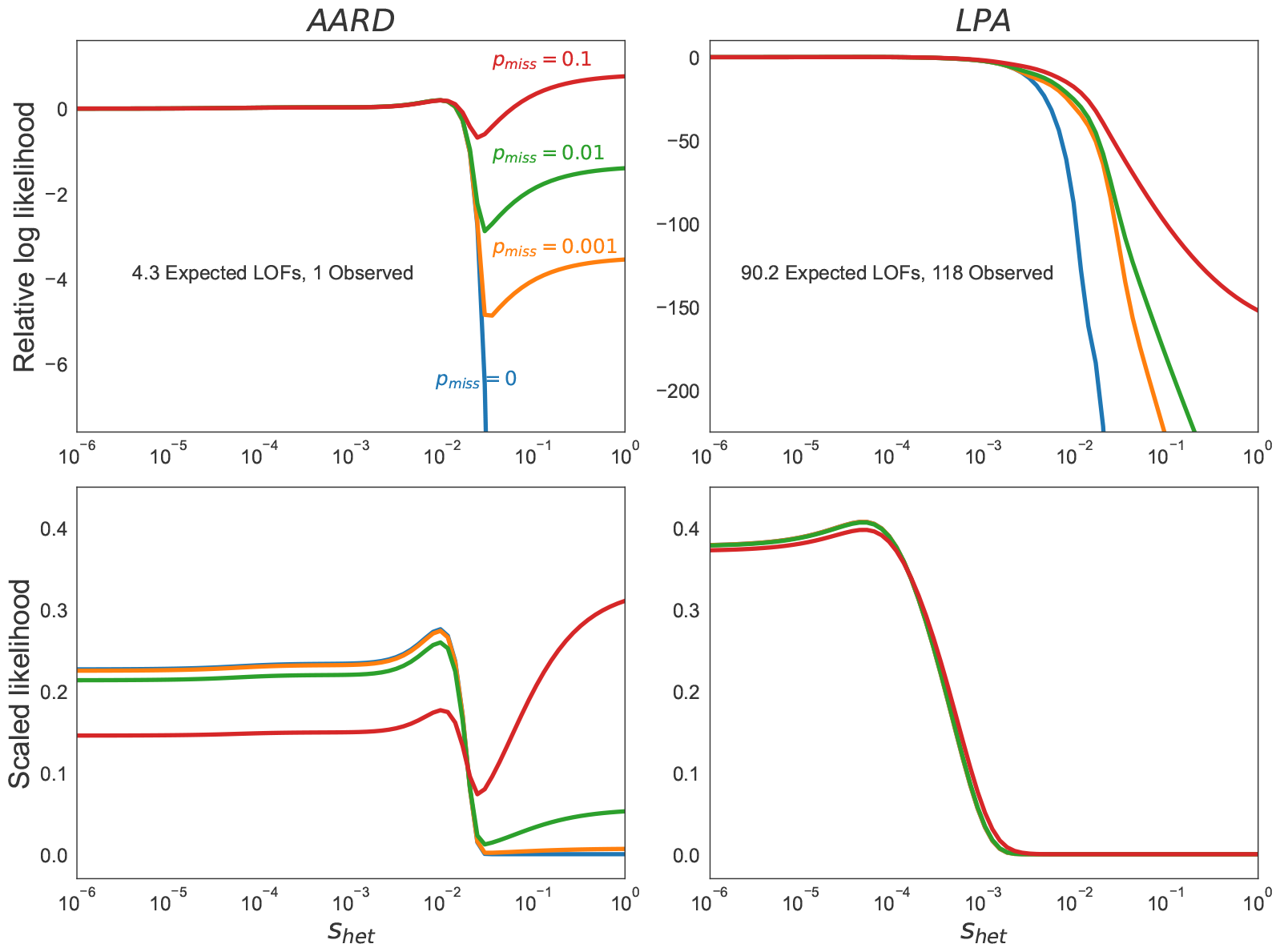
Likelihoods for example genes. Relative log likelihoods (top row) or scaled likelihoods (bottom row) for representative genes AARD (left column) and LPA (right column) for different misannotation probabilities, p_miss_. Here we set p_miss_ to be the same regardless of the type of LOF variant. AARD is a representative example of a gene with few expected LOFs, while LPA is a representative example of a gene with many expected LOFs.

**Supplementary Figure 4:**
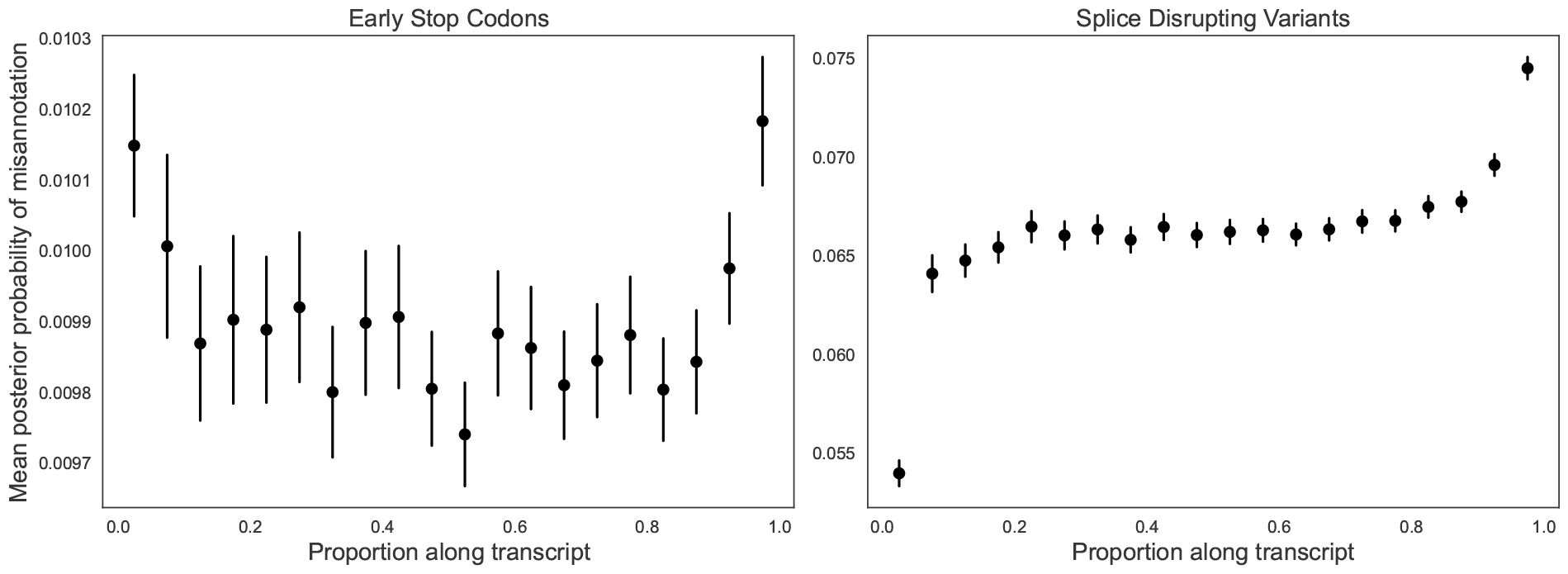
Mean posterior probabilities of misannotation along canonical transcripts. Each potential LOF was mapped to its position within the transcript normalized to the total transcript length. Variants within the first 5% of the transcript were binned, and then the next 5% of the transcript, and so on. In the plot, points represent the mean of the inferred posterior probabilities within each bin, and lines correspond to ±1.96 standard errors of the mean estimate.

**Supplementary Figure 5:**
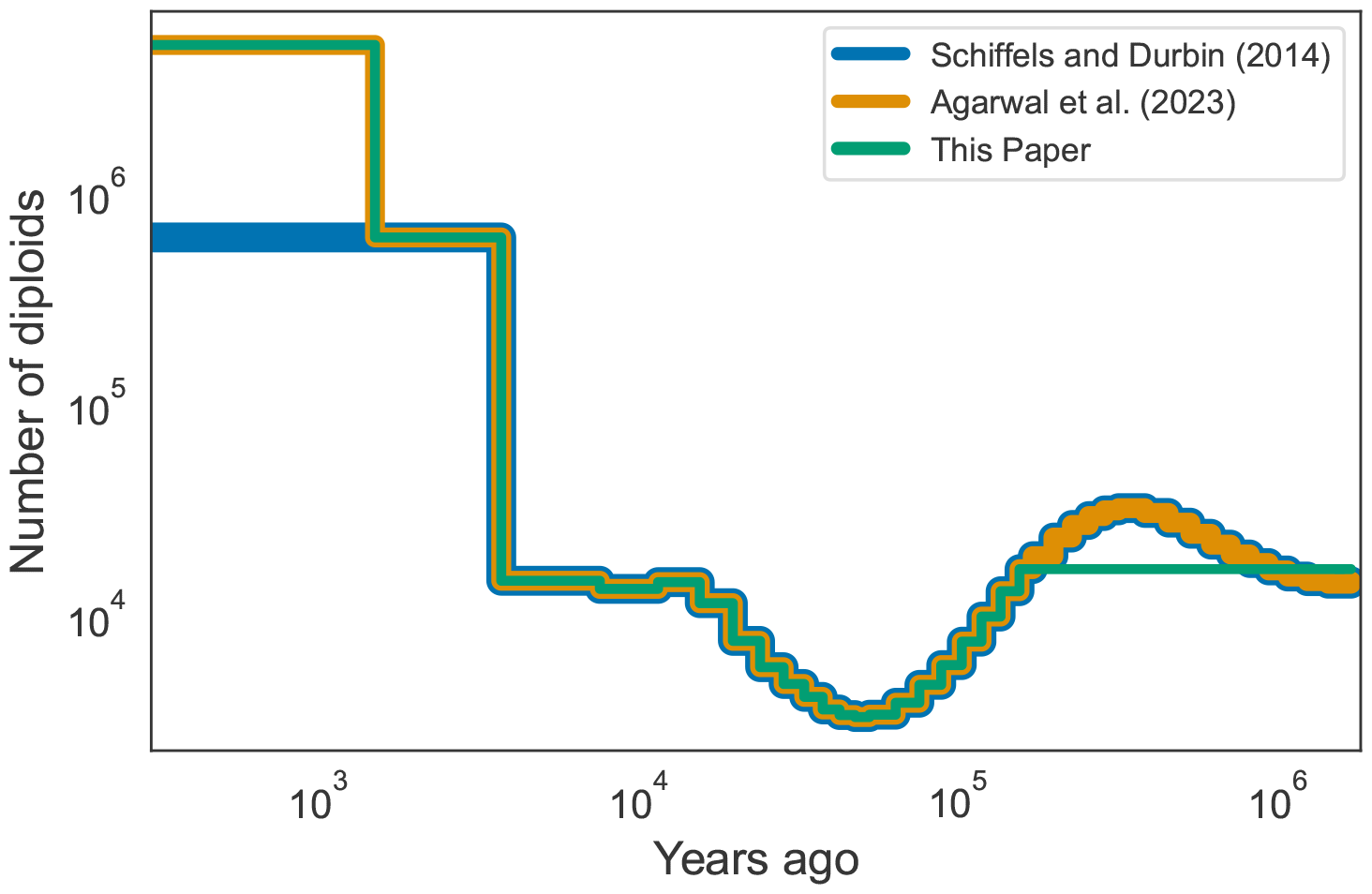
Comparison of CEU demographies. CEU demography inferred by Schiffels and Durbin [81], modified by Agarwal and colleagues [4], and further modified for this paper.

**Supplementary Figure 6:**
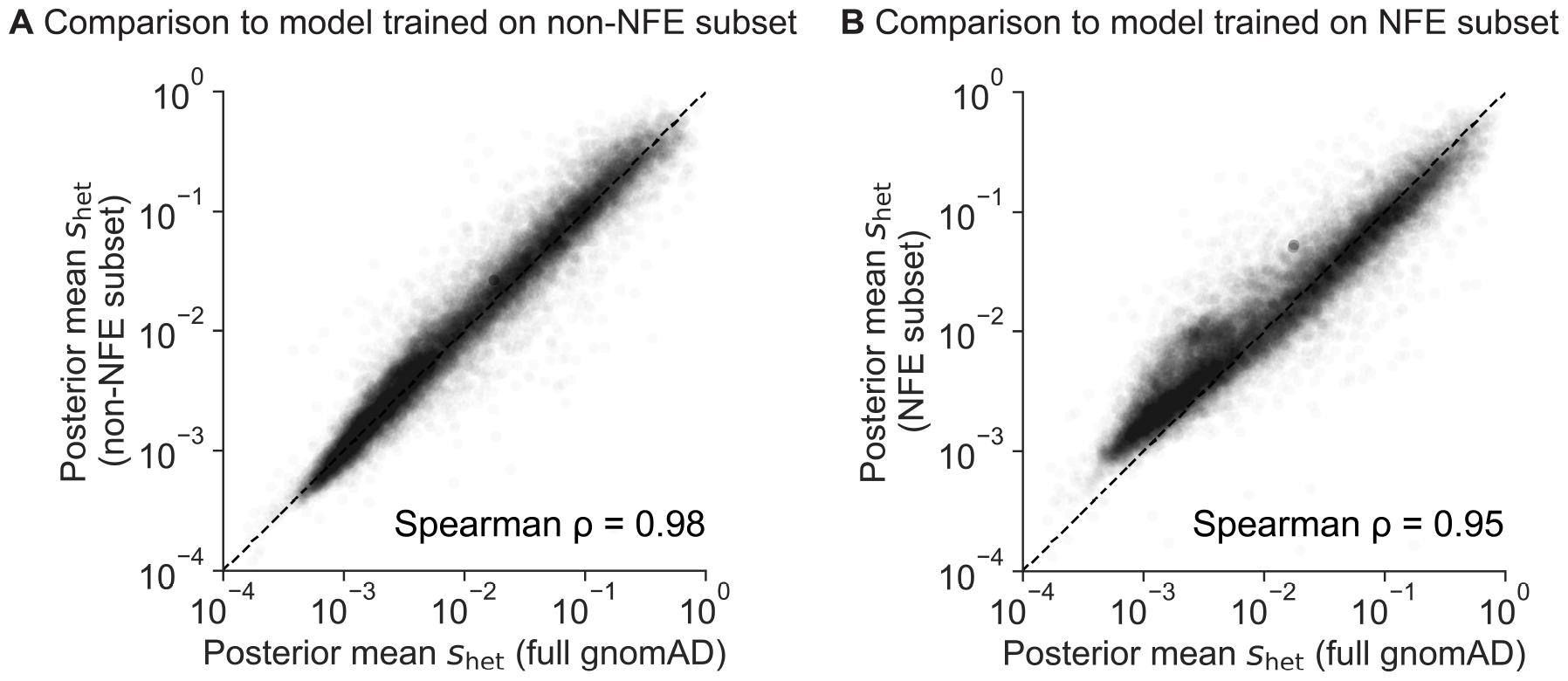
Comparison of s_het_ estimates from models trained on subsets of gnomAD. **A)** Scatterplot of posterior mean s_het_ estimated from a model trained with non-NFE individuals (y-axis) against s_het_ estimated from the full model (x-axis). NFE = Non-Finnish European. This subset consists of ∼56, 000 individuals, or ∼45% of the total dataset. **B)** Scatterplot of posterior mean s_het_ estimated from a model trained with NFE individuals (y-axis) against s_het_ estimated from the full model (x-axis). This subset consists of ∼67, 000 individuals, or ∼55% of the total dataset.

**Supplementary Figure 7:**
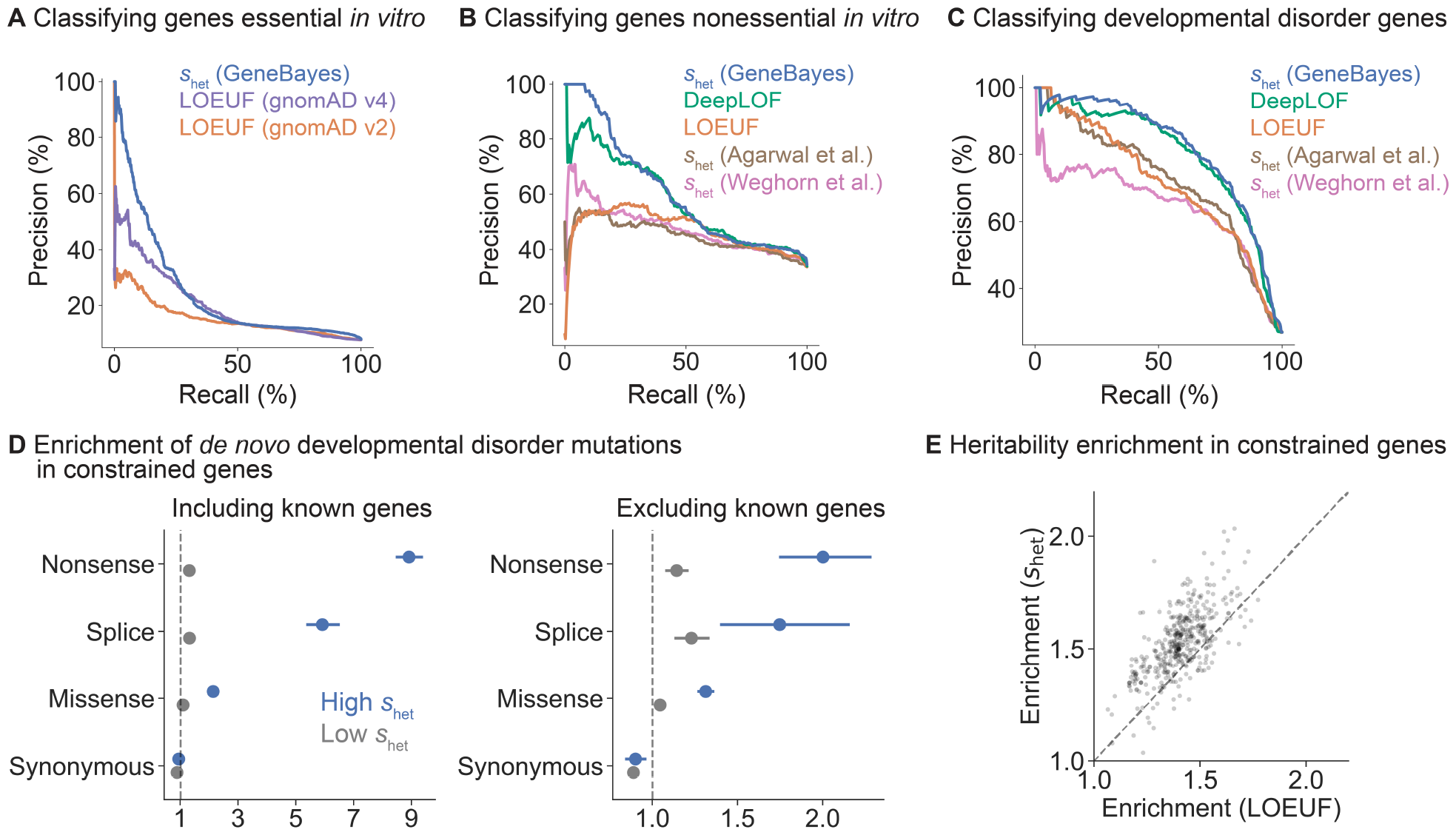
Additional validation analyses. **A)** Precision-recall curves comparing the performance of s_het_ estimates from GeneBayes against LOEUF from gnomAD v4.0.0 (∼731k exomes) or LOEUF from gnomAD v2.1.1 (∼125k exomes) in classifying essential genes. **B)** Precision-recall curves comparing the performance of s_het_ estimates from GeneBayes against other constraint metrics in classifying non-essential genes. **C)** Precision-recall curves comparing the performance of s_het_ against other constraint metrics in classifying developmental disorder genes. **D)** Enrichment of de novo mutations in patients with developmental disorders, calculated as the observed number of mutations over the expected number under a null mutational model. We plot the enrichment of synonymous, missense, splice, and nonsense variants in the 10% of genes considered most constrained by s_het_ (blue) and the enrichment of these variants in all other genes (gray), including (left) and excluding (right) known developmental disorder genes. Bars represent 95% confidence intervals. **E)** Scatterplot of the enrichment of common variant heritability in the 10% of genes considered most constrained by s_het_ (y-axis) or LOEUF (x-axis), normalized by the enrichment of heritability in all genes. Each point represents one trait.

**Supplementary Figure 8:**
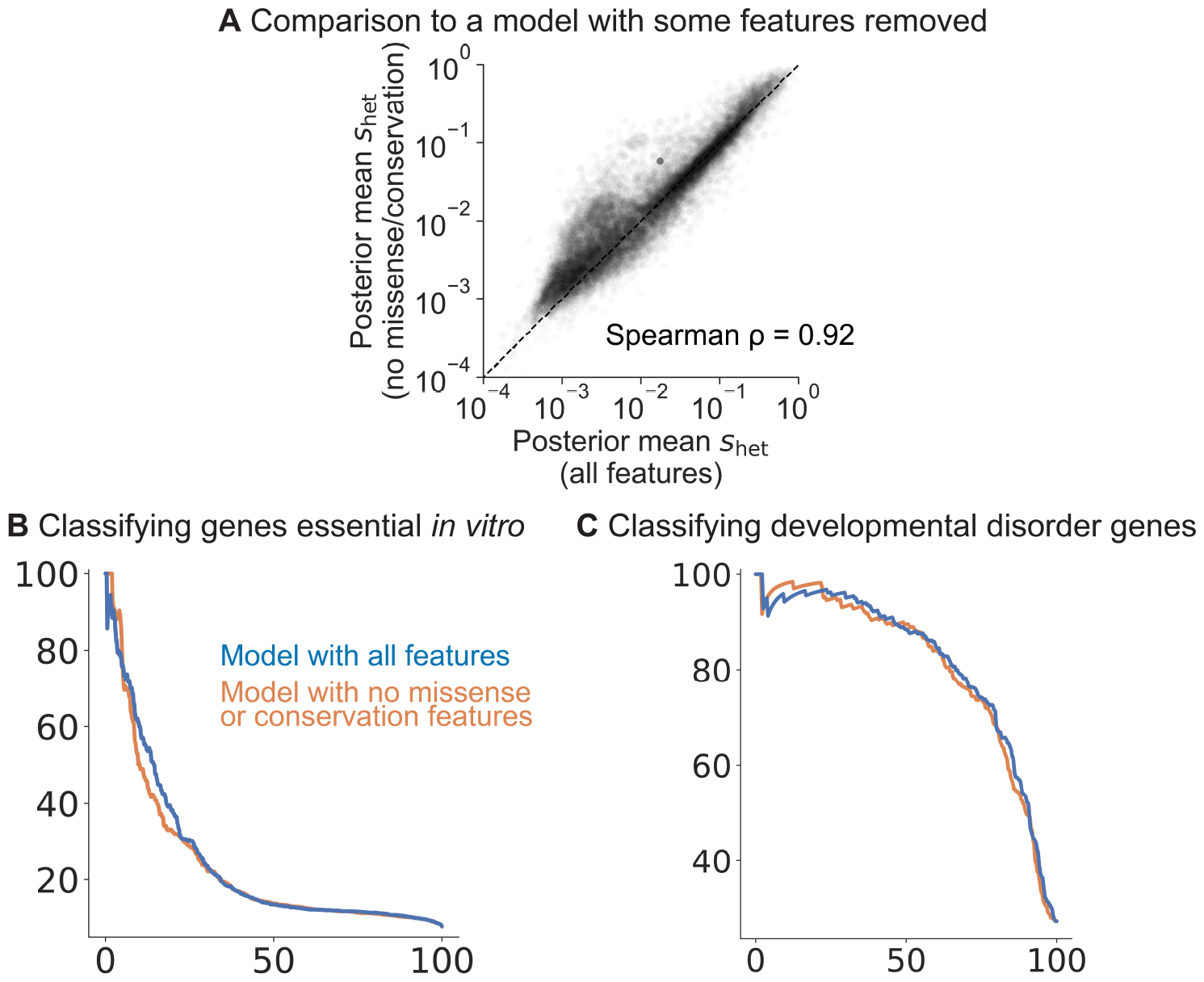
Performance of s_het_ estimates from a model with some features removed. **A)** Scatterplot of posterior mean s_het_ estimated from a model trained without missense constraint or cross-species conservation features (y-axis) against s_het_ estimated from the full model (x-axis). **B)** Precision-recall curves comparing the performance of s_het_ estimated from the full model (blue) and from the model without missense/conservation features (orange) in classifying essential genes. **C)** Precision-recall curves comparing the performance of s_het_ estimated from the two models in classifying developmental disorder genes.

**Supplementary Figure 9:**
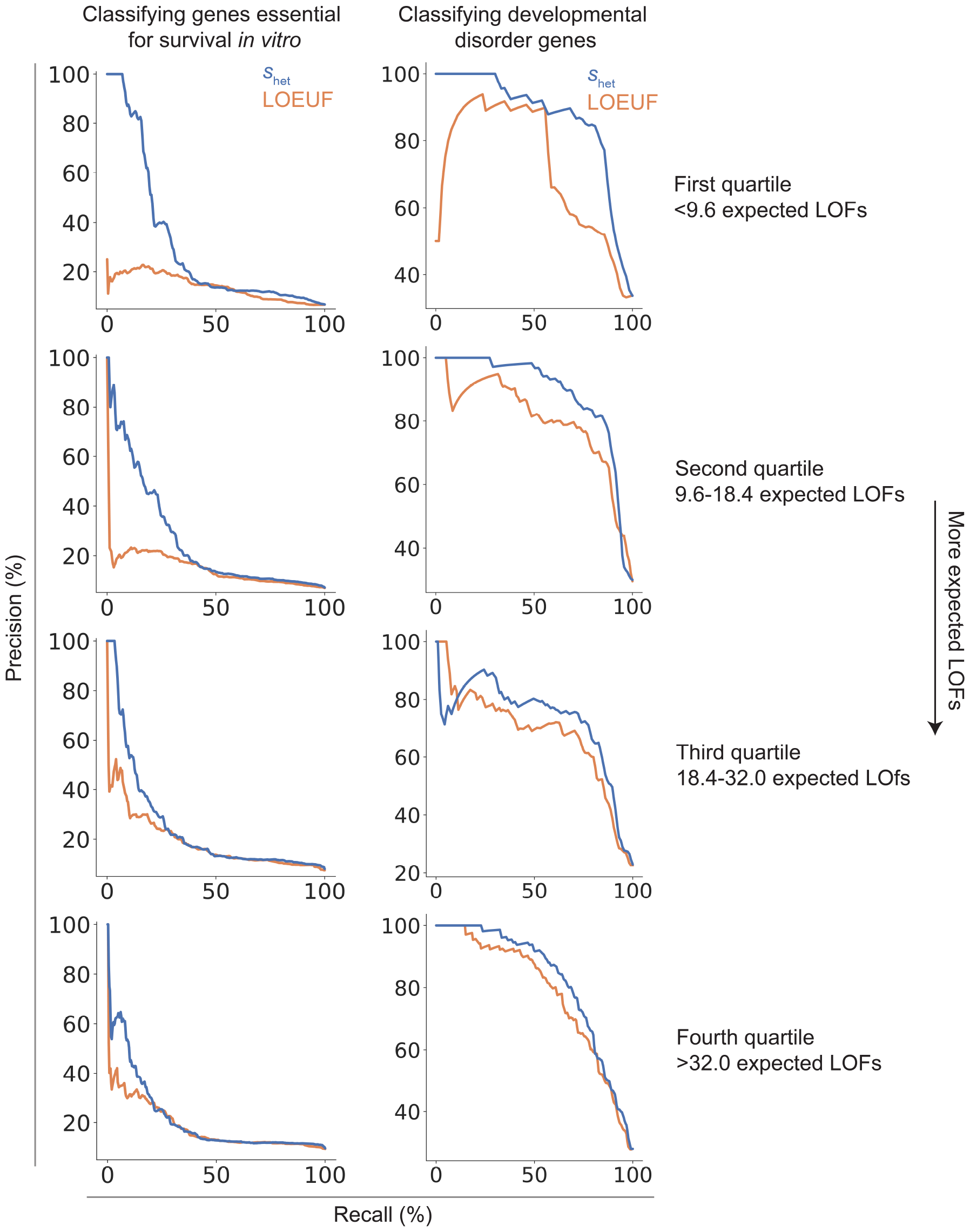
Performance of s_het_ and LOEUF for genes with differing numbers of expected LOFs. Left: Precision-recall curves comparing the performance of s_het_ against LOEUF in classifying essential genes for groups of genes binned by their expected number of LOFs. Right: Precision-recall curves comparing the performance of s_het_ against LOEUF in classifying developmental disorder genes for binned genes.

**Supplementary Figure 10:**
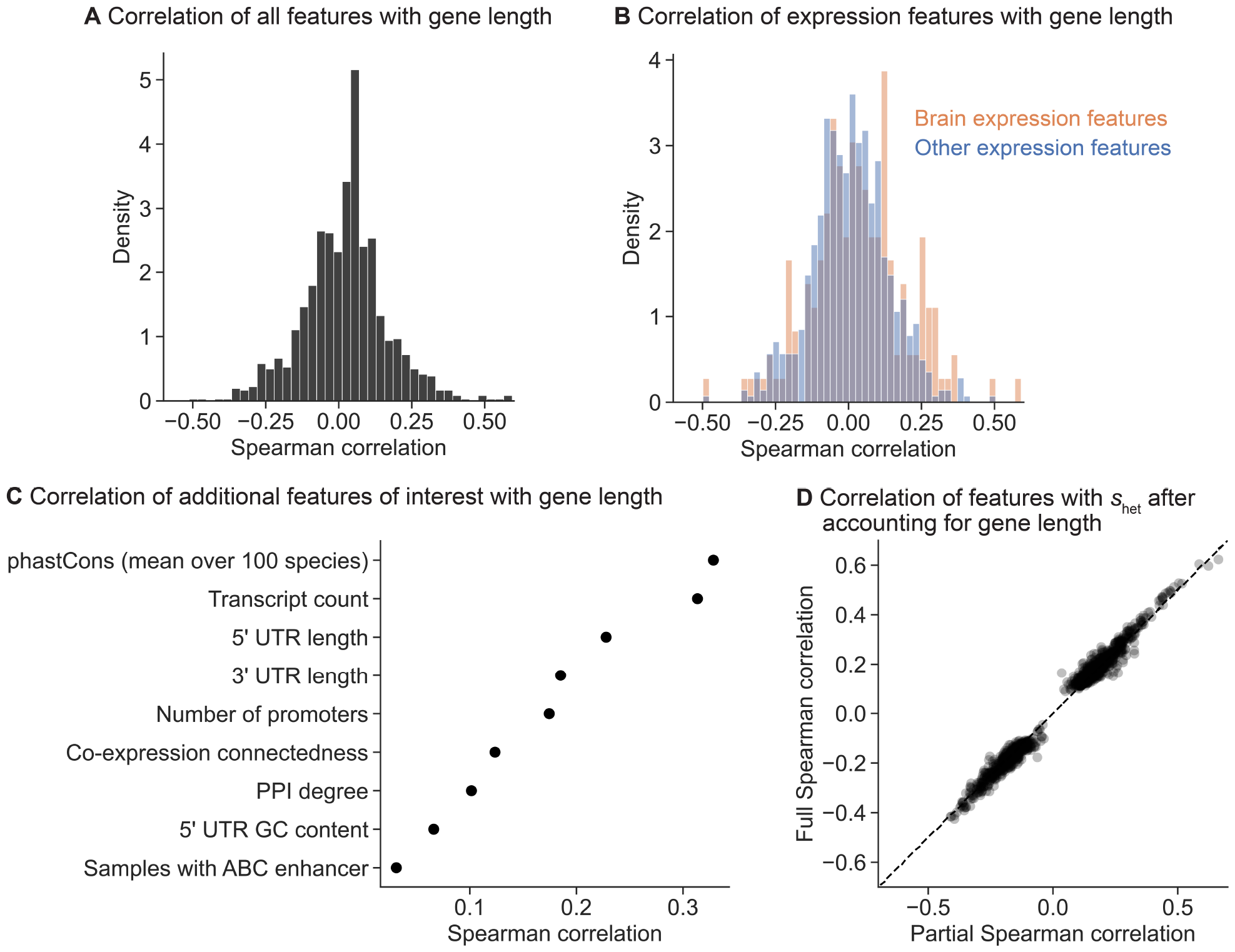
Correlation of gene features with gene length. **A)** Histogram of the Spearman ρ between gene features and coding sequence (CDS) length. **B)** Histogram of the Spearman ρ between gene features and CDS length for gene expression features, colored by category. **C)** Spearman ρ between gene features and CDS length for additional features of interest. **D)** Scatterplot of the Spearman ρ between gene features and posterior mean s_het_ (y-axis) against the partial Spearman ρ (x-axis) after controlling for the effect of gene (CDS) length.

## Notes

### Competing Interest Statement

The authors have declared no competing interest.

### Summary of Updates

We have added additional analyses and released updated estimates of s_het.

